# Machine learning methods applied to classify complex diseases using genomic data

**DOI:** 10.1101/2024.03.18.585541

**Authors:** Magdalena Arnal Segura, Giorgio Bini, Anastasia Krithara, George Paliouras, Gian Gaetano Tartaglia

## Abstract

Complex diseases pose challenges in disease prediction due to their multifactorial and polygenic nature. In this work, we explored the prediction of two complex diseases, multiple sclerosis (MS) and Alzheimer’s disease (AD), using machine learning (ML) methods and genomic data from UK Biobank. Different ML methods were applied, including logistic regressions (LR), gradient boosting decision trees (GB), extremely randomized trees (ET), random forest (RF), feedforward networks (FFN), and convolutional neural networks (CNN). The primary goal of this research was to investigate the variability of ML models in classifying complex diseases based on genomic risk. LR was the most robust method across folds and diseases, whereas deep learning methods (FFN and CNN) exhibited high variability. When comparing the performance of polygenic risk scores (PRS) with ML methods, PRS consistently performed at an average level. However, PRS still offers several practical advantages over ML methods. Despite implementing feature selection techniques to exclude non-informative and correlated predictors, the performance of ML models did not improve significantly, underscoring the ability of ML methods to achieve optimal performance even in the presence of correlated features due to linkage disequilibrium. Upon applying explainability tools to extract information about the genomic features contributing most to the classification task, the results confirmed the polygenicity of MS. The prevalence of HLA gene annotations among the top genomic features on chromosome 6 aligns with their significance in the context of MS. Overall, the highest-prioritized genomic variants were identified as expression or splicing quantitative trait loci (eQTL or sQTL) located in non-coding regions within or near genes associated with the immune response and MS. In summary, this research offers deeper insights into how ML models discern genomic patterns related to complex diseases.

## INTRODUCTION

In the context of genetics, complex diseases or conditions arise from the combined effects of multiple genomic variants and genes, are influenced significantly by both the physical and the social environment, and display non-Mendelian inheritance patterns. As a result, the task of finding genomic factors that contribute to the predisposition or protection against these diseases often requires the use of large cohorts of individuals with comprehensive genomic information at high density to obtain statistically significant results.

In recent decades, population genomics has made advances in the genomic characterization of complex diseases, largely due to the improvement in genomic technologies, the increase in the available genomic data, and the emergence of genome-wide association studies (GWAS) (Uffelmann et al. 2021). While the utility of GWAS is undeniable, there are several limitations associated with the use of this technique. First, the presence of linkage disequilibrium (LD) makes it challenging to identify the exact causal variant. Researchers often use fine-mapping tools to try to address issues derived from LD and find the causal SNVs in GWAS signals (Hormozdiari et al. 2014), (Benner et al. 2016), (Kichaev et al. 2014) (G. Wang et al. 2020). However, the genomic variants associated with the disease selected by these tools are not always consistent across methods (Uffelmann et al. 2021).

Another limitation of GWAS is that associations between SNVs and the phenotype are commonly tested using linear regression models for continuous phenotypes, or logistic regression models for binary phenotypes. Therefore, while GWAS is effective in uncovering the main effects of genomic variants within LD blocks concerning a particular condition, it is less suited for detecting interactions between genomic variants influencing disease risk. To address this limitation, several statistical methods such as the multifactor dimensionality reduction (MDR) (Moore and Andrews 2015), AprioriGWAS (Zhang, Long, and Ott 2014), fpgrowth (Nasreen et al. 2014), several Bayesian methods (Chen et al. 2021), and machine learning methods (Okazaki and Ott 2022), offer the possibility to prioritize genomic variants based on interactions.

In population genomics, the most popular statistical approach used to quantify the genomic risk of individuals to develop a trait or disease is the polygenic risk score (PRS), which uses summary statistics obtained from GWAS to fit the equations. PRS has been extensively used in many studies, demonstrating its ability to extract disease risk propensity scores from diverse cohorts and conditions (Collister, Liu, and Clifton 2022). However, a limitation still exists, as PRS are not designed to detect epistatic events associated with a condition.

Alternatively, machine learning (ML) and deep learning (DL) are subfields of artificial intelligence (AI) that focus on the development of algorithms and models that enable computers to learn from and make predictions based on data. DL is a subset of ML that focuses on neural networks with multiple layers. The emergence of big data has facilitated the application of these methods, contributing to their increasing popularity in a wide range of fields, including population genomics (Lin and Ngiam 2023). ML methods calculate the importance of input features during training, and are able to detect interactions and complex patterns in the data (Ott and Park 2022). The use of Explainable AI (XAI) methods in ML aims to provide information on the most relevant features as considered by the models for making predictions (Lipton 2016). In this context, the most informative genomic features identified by different ML methods can foster the discovery of complex genomic profiles associated with these conditions.

In this work, ML and DL methods were applied for the purpose of classifying individuals with two complex diseases, multiple sclerosis (2,020 cases) and Alzheimer’s disease (3,126 cases), in comparison to non-affected controls sourced from the UK Biobank (UKB), using data from genotyping arrays.

Multiple sclerosis (MS) is a chronic autoimmune disease of the central nervous system where the immune system mistakenly attacks the protective covering of nerve fibers (myelin), leading to communication disruptions between the brain and the rest of the body. This can result in a wide range of symptoms, including fatigue, difficulty walking, numbness or tingling, and problems with coordination and balance (McGinley, Goldschmidt, and Rae-Grant 2021). There have been several published works aiming to identify individuals with MS and their associated genetic loci linked to the progression of disability in the disease by applying ML methods (Fuh-Ngwa et al. 2022) (Ghafouri-Fard et al. 2020). In addition, the detection of epistatic events among genomic variants associated with MS using ML methods have been studied in other works (Burnard et al. 2022) (Briggs and Sept 2021).

Alzheimer’s disease (AD) is a progressive neurodegenerative disorder characterized by cognitive decline, memory loss, and changes in behavior, primarily affecting older individuals. The disease is associated with the accumulation of abnormal protein deposits, including beta-amyloid plaques and tau tangles, in the brain (Ballard et al. 2011). ML classifiers have been previously used to classify AD using genotyping data. Authors in a published work (De Velasco Oriol et al. 2019) examined the application of six different ML methods, including RF, to predict the risk of AD using genomic data from the Alzheimer’s Disease Neuroimaging Initiative (ADNI) cohort. The research systematically compared various ML models and found that the best-performing models achieved around 0.72 Area Under the Receiver Operating Characteristic Curve (AUC-ROC), indicating their potential for predicting AD risk. Another study contrasted different ML methods and proposed enhancing prediction by adding markers from misclassified samples (Romero-Rosales et al. 2020).

Regarding the application of DL methods to AD, authors in a study tried to address the problem of the limited population representativity by creating a DL-based framework designed to enhance the accuracy of genetic risk prediction by incorporating data from diverse populations (Gyawali et al. 2023). In addition XAI tools applied to DL models in AD have been proposed in several studies (Jemimah and AlShehhi 2023), (Chandrashekar et al. 2023), (Vivek et al. 2023), (Chang et al. 2020) and (Lundberg et al. 2023).

Overall, the application of ML methods to identify individuals with complex diseases based on genomic data has gained popularity in recent years. However, a limitation is that the published works on this topic are relatively recent, and there are not yet many studies evaluating the robustness of these methods. Additionally, some of the studies referenced in the previous lines used relatively small sample sizes in the analysis (fewer than 1,000 cases), which makes it challenging to draw generalizable conclusions.

The primary objective of this study is to assess the variability and robustness of ML techniques in predicting complex diseases using genomic data. This is relevant because the results of ML methods may vary depending on the data, the design of the model, the strategy used for training, and the hyperparameter space, among other reasons. In addition, genomic variants inherently exhibit correlation due to LD, which may negatively impact model performance. In this work, we also compared the performance of ML models to the PRS. The secondary goal of this study is to apply XAI tools to the ML models to extract information about the prioritized features that contributed the most in the classification task, pointing to potential predisposing or protective genomic variants in the disease.

After exploring these aspects, we aim to provide insights into the considerations to take into account when using ML methods for the classification of complex diseases based on genomic data.

## RESULTS

### Performance of the models

The ML methods employed in this work were logistic regression (LR), three tree-based ensemble ML methods, including Gradient-Boosted Decision Trees (GB) (Friedman 2001), Random Forest (RF) (Breiman 2001), and Extremely Randomized Trees (ET) (Geurts, Ernst, and Wehenkel 2006), and two DL methods including Feedforward networks (FFN) and Convolutional Neural Networks (CNN) (see supplementary material for details on the ML methods). The predictors included in the ML models were genomic variants associated with the disease reported in curated databases.

Table 1 (a) and (b) show the evaluation metrics for models constructed with MS and AD, respectively. The mean and standard deviation of different evaluation metrics across the five folds in the outer loop of the nested CV are provided. The mean performance scores for both diseases typically ranged around 0.6 and 0.7, with few exceptions. Notably, FFN and CNN methods exhibited the least stable performance, as evidenced by the highest standard deviation across folds. In the case of AD, GB method performed similarly to CNN and FFN with low mean performances and high standard deviation, while GB demonstrated relatively good performance in MS.

**Table 1.** comprises two independent tables showing the mean and standard deviation of evaluation metric values across the five folds in the outer loop of the nested CV. The evaluation metrics represented in the table include balanced accuracy, specificity, sensitivity, and AUC-ROC. For each column, the color scale ranges from darker to lighter, indicating better to worse performance, respectively. (a) presents results corresponding to MS, while (b) presents results corresponding to AD.

|  | accuracy mean | accuracy sd | specificity mean | specificity sd | sensitivity mean | sensitivity sd | AUC-ROC mean | AUC-ROC sd |
| --- | --- | --- | --- | --- | --- | --- | --- | --- |
| **GB** | 0.628 | 0.007 | 0.635 | 0.005 | 0.622 | 0.017 | 0.670 | 0.009 |
| **ET** | 0.625 | 0.006 | 0.660 | 0.014 | 0.590 | 0.022 | 0.660 | 0.006 |
| **RF** | 0.612 | 0.008 | 0.657 | 0.011 | 0.567 | 0.022 | 0.655 | 0.011 |
| **LR** | 0.635 | 0.005 | 0.635 | 0.008 | 0.634 | 0.010 | 0.690 | 0.009 |
| **FFN** | 0.629 | 0.014 | 0.599 | 0.059 | 0.660 | 0.075 | 0.674 | 0.012 |
| **CNN** | 0.619 | 0.011 | 0.652 | 0.058 | 0.587 | 0.067 | 0.654 | 0.018 |

Table 1 comprises two independent tables showing the mean and standard deviation of evaluation metric values across the five folds in the outer loop of the nested CV.
|  | accuracy mean | accuracy sd | specificity mean | specificity sd | sensitivity mean | sensitivity sd | AUC-ROC mean | AUC-ROC sd |
| --- | --- | --- | --- | --- | --- | --- | --- | --- |
| **GB** | 0.637 | 0.021 | 0.651 | 0.022 | 0.623 | 0.034 | 0.671 | 0.021 |
| **ET** | 0.675 | 0.013 | 0.723 | 0.006 | 0.627 | 0.031 | 0.708 | 0.011 |
| **RF** | 0.681 | 0.011 | 0.723 | 0.007 | 0.639 | 0.026 | 0.709 | 0.014 |
| **LR** | 0.674 | 0.010 | 0.693 | 0.005 | 0.655 | 0.024 | 0.715 | 0.012 |
| **FFN** | 0.645 | 0.018 | 0.693 | 0.068 | 0.598 | 0.072 | 0.694 | 0.026 |
| **CNN** | 0.629 | 0.024 | 0.665 | 0.043 | 0.594 | 0.034 | 0.667 | 0.024 |

Conversely, ET and RF were among the best methods in the context of AD and showed worse performance in MS. These findings emphasize the variability in the performance of DL and tree-based methods when tested across various folds and diseases. In both diseases, LR method exhibited low values of standard deviation and displayed consistent results across various evaluation metrics.

External validation datasets with cohorts from the International Multiple Sclerosis Genetics Consortium (IMSGC) and the Alzheimer’s Disease Neuroimaging Initiative (ADNI) were used to evaluate the model’s generalization performance for MS and AD models, respectively (see supplementary material). Sensitivity or balanced accuracy did not worsed compared to the previously reported results in UKB, supporting the model’s ability to generalize across diverse populations, and dismissing concerns about potential overfitting.

### Comparison of machine learning methods with polygenic risk score

The performance of ML methods was compared with PRS. The same division of samples into folds was used for the PRS calculation and ML methods, comparing the same individuals across the 5 folds. We followed the standard practice for PRS calculations, which is to use all the genomic variants that have successfully passed the quality filters present in both target and base data. Therefore, the number of predictors included in PRS models was higher than the one in ML methods (see supplementary tables).

The ML models employed in this work, which are classifiers, return the probabilities assigned to each subject. Instead, PRS return risk scores as continuous values without any fixed range. To convert PRS scores into predicted classes, the regular practice is to set a cut-off using percentiles. In this work, we applied percentiles to the probabilities obtained with ML methods and to the values of PRS to compare their performance. The relative risk (RR) and odds ratio (OR) were calculated by considering individuals with the highest values of PRS and ML probabilities (upper 99th percentile) as predicted positives. Values of RR and OR higher than one indicated a higher proportion of individuals with the disease in the upper 99^th^ percentile compared to the rest of individuals in lower percentiles.

The mean and standard deviation of RR and OR across the five folds are provided in Table 2 for MS and AD. For ML methods, results are similar as the ones obtained for the general evaluation metrics presented in Table 1, with LR doing relatively well across diseases. For MS, PRS was among the top three best methods after LR and FFN as shown in Table 2 (a). Consequently, LR and FFN proved to be more effective at stratifying the risk of MS, even when using a lower number of predictors in the models with respect to PRS. In AD, PRS had average performance when compared with the other ML methods, as shown in Table 2 (b).

**Table 2.** comprises two tables showing the mean and standard deviation of the relative risk (RR) and odds ratio (OR) across the samples used in the five folds. The formulas used in the calculation of RR and OR are provided in the Methods section 5.4. RR and OR were calculated considering as positives the samples ranked within the top 99th percentile with the best scores or probabilities. In the case of ML methods, the cutoff of probability 0.5, which is the default in these methods, was also considered to define positives and calculate the RR and OR and is represented in additional columns. Results for MS and AD are presented in tables (a) and (b), respectively. For each column, the color scale ranges from darker to lighter, indicating better to worse performance, respectively.

|  | RR mean<br>99th percentile | RR sd<br>99th percentile | OR mean<br>99th percentile | OR sd<br>99th percentile | RR mean<br>ML classification | RR sd<br>ML classification | OR mean<br>ML classification | OR sd<br>ML classification |
| --- | --- | --- | --- | --- | --- | --- | --- | --- |
| GB | 3.598 | 1.575 | 3.901 | 1.822 | 2.790 | 0.162 | 2.868 | 0.170 |
| ET | 3.886 | 1.653 | 4.247 | 2.004 | 2.719 | 0.119 | 2.795 | 0.125 |
| RF | 3.341 | 1.687 | 3.610 | 1.915 | 2.456 | 0.159 | 2.517 | 0.168 |
| LR | 5.509 | 1.390 | 6.232 | 1.782 | 2.935 | 0.119 | 3.021 | 0.126 |
| FFN | 4.107 | 0.630 | 4.455 | 0.752 | 2.908 | 0.426 | 2.988 | 0.444 |
| CNN | 2.827 | 1.060 | 2.984 | 1.219 | 2.641 | 0.269 | 2.712 | 0.284 |
| PRS | 4.002 | 0.729 | 4.333 | 0.864 | --- | --- | --- | --- |

Table 2 comprises two tables showing the mean and standard deviation of the relative risk (RR) and odds ratio (OR) across the samples used in the five folds.
|  | RR mean<br>99th percentile | RR sd<br>99th percentile | OR mean<br>99th percentile | OR sd<br>99th percentile | RR mean<br>ML classification | RR sd<br>ML classification | OR mean<br>ML classification | OR sd<br>ML classification |
| --- | --- | --- | --- | --- | --- | --- | --- | --- |
| GB | 4.313 | 0.823 | 4.795 | 1.037 | 3.013 | 0.564 | 3.128 | 0.609 |
| ET | 6.643 | 1.315 | 8.031 | 1.914 | 4.183 | 0.426 | 4.408 | 0.462 |
| RF | 6.837 | 0.719 | 8.273 | 1.044 | 4.391 | 0.415 | 4.636 | 0.453 |
| LR | 6.882 | 0.651 | 8.336 | 0.969 | 4.094 | 0.368 | 4.300 | 0.398 |
| FFN | 5.906 | 1.696 | 7.011 | 2.528 | 3.350 | 0.611 | 3.502 | 0.673 |
| CNN | 4.788 | 1.798 | 5.473 | 2.487 | 2.862 | 0.608 | 2.971 | 0.661 |
| PRS | 5.683 | 1.098 | 6.644 | 1.523 | --- | --- | --- | --- |

As previously exposed, PRS do not provide probabilities but instead offer risk scores associated with the disease, which are used to identify individuals at high risk and low risk. Instead, ML methods were employed as classifiers in the previous section of this work, applying a cut-off of probability 0.5, which is the default setting used to classify samples as positives (greater than or equal to 0.5) or negatives (lower than 0.5). The values of RR and OR considering the default cut-off used in the ML classification are provided in Table 2 as well. In MS and AD, results of RR and OR based on the ML classification are lower with respect to considering the 99^th^ percentile of samples with the highest probability as predicted positives. This fact suggests that the use of a cut-off of probability 0.5 in ML methods reduces the proportion of true positives over the false positives with respect to using the 99^th^ percentile cut-off. Related to this, FFN demonstrated the lowest standard deviation across folds when considering the top 99^th^ percentile of samples with the highest probabilities as predicted positives in MS. The 99^th^ percentile corresponded to the values of probability higher than or equal to 0.89 obtained with the FFN models. Conversely, FFN exhibited the highest standard deviation when using the probability cut-off of 0.5. These results suggest that FFN models applied to MS exhibited greater robustness in stratifying the positive class when a higher cut-off was set to split the classes.

We investigated whether individuals predicted as positives over the 99^th^ percentile or predicted as negatives under the 50^th^ percentile by the PRS models were consistently classified as positives and negatives across the six ML methods (Figure 1). In MS, 77% of cases predicted as positives by PRS were also classified as positives by the six ML methods, as indicated by the yellow bar in Figure 1 (a). Conversely, 70% of the samples classified as false positives (FP) in PRS were also misclassified by the six ML methods, as indicated by the dark blue bar in Figure 1 (b).

**Figure 1.**
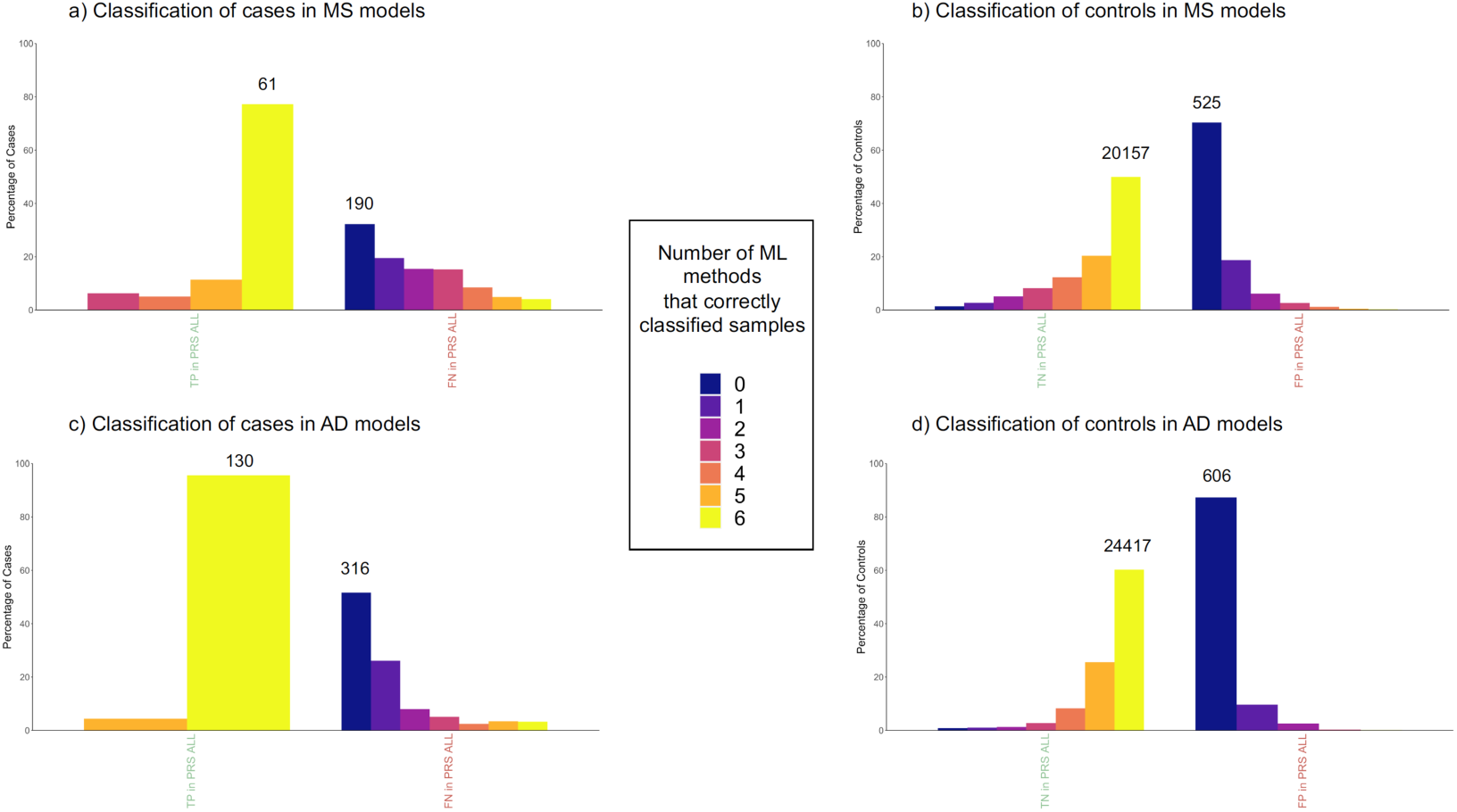
shows the percentage of MS and controls that were correctly classified by 0 to 6 ML methods in comparison with samples correctly classified (TP or TN labeled in green) or incorrectly classified (FN and FP labeled in red) by PRS models. The total number of samples is indicated above the bars for the groups with the highest percentage in each comparison. Plots (a) and (b) show the classification of cases and controls in MS models, respectively. Plots (c) and (d) show the classification of cases and controls in AD models, respectively.

In AD, the percentage of true positives (TP) in PRS with full agreement across ML methods was 96%, as indicated in the yellow bar of Figure 1 (c), while the percentage of FP in PRS with full agreement across ML methods was 87%, as represented with the dark blue bar of Figure 1 (d).

Overall, the results presented in this section suggest that, with the evaluation based on percentiles, the performance of PRS is similar to that of the ML models. Additionally, the results in Figure 1 suggest that PRS and ML models demonstrate consistent classification results, with similarities in the classification of specific individuals. The strengths and weaknesses of ML methods and PRS will be further elaborated upon in the discussion.

### Implementation of feature selection techniques

In this study we employed curated databases of disease-related variants to select the predictors for the ML models. However, these databases contain genomic variants from diverse studies conducted in various human populations, some of which may not be informative in the UKB cohort. Furthermore, certain genomic variants used as features in models are highly correlated due to LD, with a potential negative impact in the performance of models. To address this, we used feature selection techniques such as recursive feature elimination (RFE) and recursive feature elimination with cross-validation (RFECV) aiming to identify a subset of features with the potential to enhance model performance. Because of the considerable variability observed across folds in DL methods, which could potentially compromise the robustness of the comparisons, together with the fact that in DL, the direct application of feature selection techniques is less common (Alzubaidi et al. 2021), these tools were not applied to DL methods.

As shown in Figure 2, there are no significant differences in sensitivity or specificity when comparing models after applying RFECV and RFE with the original models. However, reducing the number of features in the models with RFE and RFECV had the effect of decreasing the number of correlated features due to LD. This trend is represented in supplementary material. Therefore, the results suggest that the presence of correlation among the genomic variants did not significantly impact the model’s performance.

**Figure 2.**
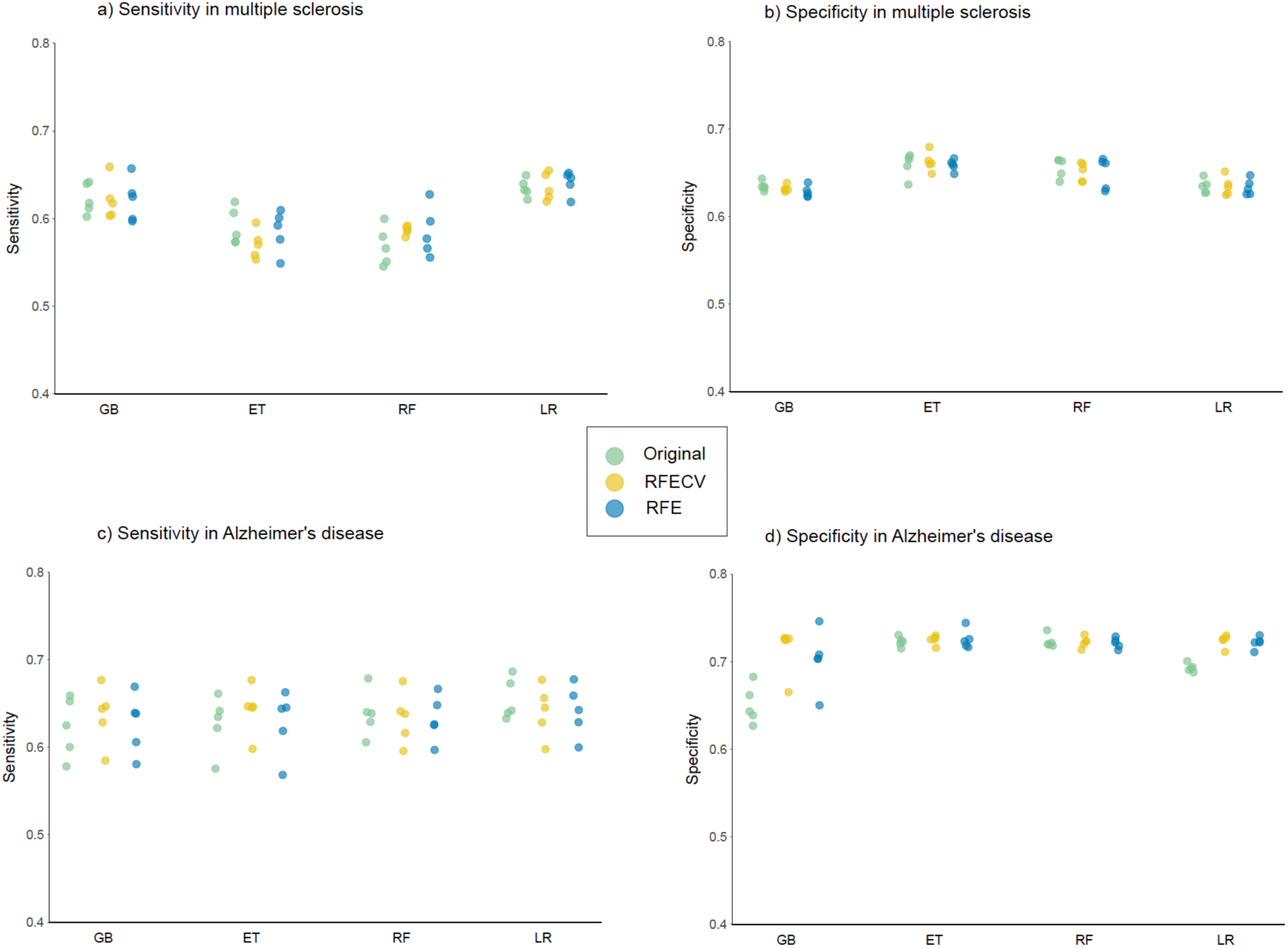
shows dot plots with the values of sensitivity and specificity in the original models, and the models after feature selection with RFECV and RFE. Sensitivity and specificity in MS are represented in plots (a) and (b), respectively. Sensitivity and specificity in AD are represented in plots (c) and (d), respectively.

In the case of MS, 74% to 88% of samples were classified with the same class using the features in the original models, RFE and RFECV, as shown in Table 3 (a). Notably, the value is particularly high for AD (see Table 3(b)) where, except for the GB method, around 93% of samples were classified with the same class using different sets of features. These results indicate that the original models, RFE and RFECV lead to the same prediction for the majority of individuals.

**Table 3.** comprises two tables showing, for each ML method, from left to right, the percentage of samples that were classified with the same class in the original models and models after feature selection, the distinct methods used for feature selection, the number of features in folds from one to five after feature selection, and the number of features selected from one to five times in different folds. In (a), the table corresponds to MS. In (b), the table corresponds to AD, and the values in red correspond to the SNV rs429358, selected across all folds and methods in AD.

|  | % samples with the same prediction in original, RFE and RFECV | Method | Features fold1 | Features fold2 | Features fold3 | Features fold4 | Features fold5 | n Features 1 time | n Features 2 times | n Features 3 times | n Features 4 times | n Features 5 times |
| --- | --- | --- | --- | --- | --- | --- | --- | --- | --- | --- | --- | --- |
| GB | 78.31 | RFE | 150 | 250 | 250 | 250 | 200 | 54 | 33 | 40 | 65 | 120 |
|  |  | RFECV | 314 | 138 | 252 | 321 | 341 | 19 | 29 | 56 | 124 | 125 |
| ET | 75.76 | RFE | 200 | 200 | 150 | 50 | 200 | 42 | 40 | 52 | 88 | 34 |
|  |  | RFECV | 86 | 316 | 274 | 175 | 277 | 33 | 29 | 87 | 109 | 68 |
| RF | 74.10 | RFE | 100 | 250 | 100 | 100 | 200 | 62 | 66 | 38 | 28 | 66 |
|  |  | RFECV | 225 | 103 | 216 | 79 | 251 | 30 | 49 | 85 | 44 | 63 |
| LR | 88.35 | RFE | 200 | 200 | 20 | 250 | 50 | 89 | 127 | 76 | 21 | 13 |
|  |  | RFECV | 205 | 140 | 25 | 231 | 39 | 102 | 128 | 50 | 18 | 12 |

Table 3 comprises two tables showing, for each ML method, from left to right, the percentage of samples that were classified with the same class in the original models and models after feature selection, the distinct methods used for feature selection, the number of features in folds from one to five after feature selection, and the number of features selected from one to five times in different folds.
|  | % samples with the same prediction in original, RFE and RFECV | Method | Features fold1 | Features fold2 | Features fold3 | Features fold4 | Features fold5 | n Features 1 time | n Features 2 times | n Features 3 times | n Features 4 times | n Features 5 times |
| --- | --- | --- | --- | --- | --- | --- | --- | --- | --- | --- | --- | --- |
| GB | 74.68 | RFE | 5 | 150 | 5 | 5 | 5 | 140 | 9 | 1 | 1 | 1 |
|  |  | RFECV | 1 | 29 | 1 | 1 | 1 | 28 | 0 | 0 | 0 | 1 |
| ET | 93.02 | RFE | 150 | 5 | 100 | 100 | 5 | 34 | 35 | 75 | 4 | 3 |
|  |  | RFECV | 32 | 1 | 1 | 2 | 1 | 30 | 1 | 0 | 0 | 1 |
| RF | 94.38 | RFE | 150 | 5 | 100 | 100 | 100 | 32 | 16 | 22 | 75 | 5 |
|  |  | RFECV | 139 | 3 | 130 | 124 | 99 | 7 | 11 | 29 | 91 | 3 |
| LR | 92.23 | RFE | 20 | 5 | 20 | 50 | 20 | 29 | 21 | 10 | 1 | 2 |
|  |  | RFECV | 7 | 1 | 7 | 51 | 3 | 44 | 4 | 4 | 0 | 1 |

With few exceptions, in MS (Table 3(a)), the number of features selected by RFECV and RFE in each fold exceeded that in AD (Table 3(b)). In fact, for AD, RFECV selected only one SNV in eight folds (Table 3(b) highlighted in red). As noted in the column labelled “n features 5 times”, 12 to 125 features were consistently selected across the five folds in MS (Table 3(a)), while AD had only 1 to 5 features selected (Table 3(b).

In AD, the missense variant rs429358 in the *Apolipoprotein E* (*APOE*) gene was the one consistently chosen across all folds using various feature selection techniques and ML methods, and it was the only feature selected in the eight different folds following RFECV selection (Table 3(b) highlighted in red). rs429358 is a SNV with the minor and major alleles being (C) and (T) respectively. The fact that RFECV proposed models with the variant rs429358 alone, without any noticeable impact on model performance, suggests that the majority of predictions were entirely influenced by this variant in the original models.

To support this assumption, Table 4 represents the allele frequency (AF) and the percentage of individuals with the rs429358 (C) allele, present in either the heterozygous or homozygous form, across different groups. In individuals with AD, the presence of rs429358 (C) was more than the double than in controls. AD individuals that were classified as true positives across all ML methods were 64% (C;T) and 36% (C;C), and with the exception of three AD subjects, all of them had at least one copy of the rs429358 (C) allele. In contrast, when looking at the AD individuals that were consistently classified as AD across ML and PRS, all of them had at least one rs429358 (C) allele, 91% of them had (C;C) alleles, while 9% had (C;T) alleles. Therefore, individuals with the highest risk of developing the disease according to PRS and ML methods seem to match those having the (C;C) alleles, followed by individuals with (C;T) alleles, in an additive risk pattern. These results demonstrate that the models constructed for AD predominantly relied on a single SNV, in this case rs429358, and that the high consistency observed in the classification of individuals across different methods for this disease is primarily attributed to this variant.

**Table 4.** shows, in columns from left to right, the allele frequency of the rs429358 (C) minor allele, the percentage of individuals with the heterozygous form of the allele (C;T), and the percentage of individuals with the two copies of the minor allele (C;C). In rows from top to bottom, there are controls, individuals with AD, AD that were correctly classified by the six ML methods, and AD that were correctly classified by the six ML methods and PRS. For each column, the color scale ranges from darker to lighter, indicating higher to lower values, respectively.

|  | AF (C) | % (C;T) | % (C;C) |
| --- | --- | --- | --- |
| **Controls** | 0.147 | 25.34 | 1.99 |
| **AD** | 0.394 | 48.96 | 14.94 |
| **TP across ML** | 0.678 | 63.73 | 35.97 |
| **TP across ML and PRS** | 0.955 | 8.94 | 91.06 |

In the case of MS, the HLA variant *HLA-A*02:01* was the only genomic variant consistently selected across different folds and methods. However, we discarded the possibility that MS models relied only on this variant for making the predictions. This conclusion is supported by the fact that the number of features used in the models after feature selection ranged from 20 to 341 features, suggesting the presence of polygenicity.

### Prioritized genomic variants in multiple sclerosis

To proceed with the use of explainability tools and extract the importance of the features assigned by the models, we focused only on MS. AD was excluded from these analysis because, as demonstrated in the previous section, the classification for this disease heavily relied on a single SNV.

For each ML model, the genomic variants were ranked with ordinal numbers, with values close to one representing higher importance (supplementary tables). In total, there were 136 genomic variants that were among the top 10% with the best rank at least in one ML method, with 50 of them present on chromosome 6. From now on, these will be referred to as prioritized variants. The circos plot in Figure 3 includes a heatmap depicting the ranks and locations of all the genomic variants used as features in MS. The genomic variants that were prioritized by at least one method are annotated with the name, excluding variants in chromosome 6. Due to the high density of prioritized genomic variants in chromosome 6, this chromosome was independently represented in Figure 4, along with the names of the 10 highest-ranked variants and the pairwise LD.

**Figure 3:**
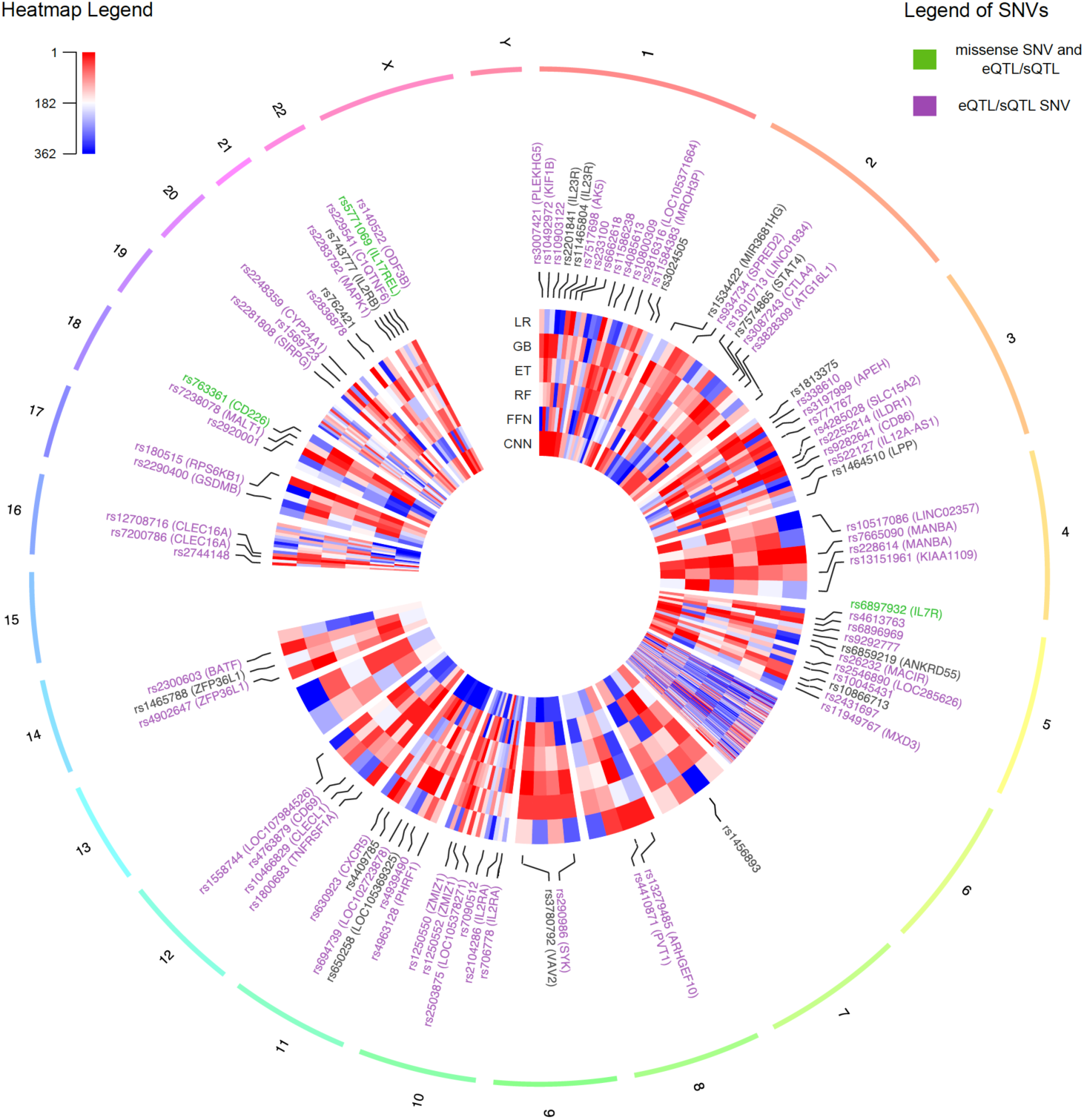
Circos plot representing all the genomic features used in the MS models distributed across the genome. The heatmap indicates the ranks of the features as assigned by each ML method, with values close to one in red indicating higher importance. The variants that were prioritized by at least one method are indicated with their names. The names of the SNVs are colored in purple if they are annotated with an eQTL or sQTL in at least one tissue in GTEx. The labels of missense SNVs with annotated QTLs are colored in green. The labels of chromosome 6 were excluded due to the high density of prioritized genomic variants in this chromosome.

**Figure 4:**
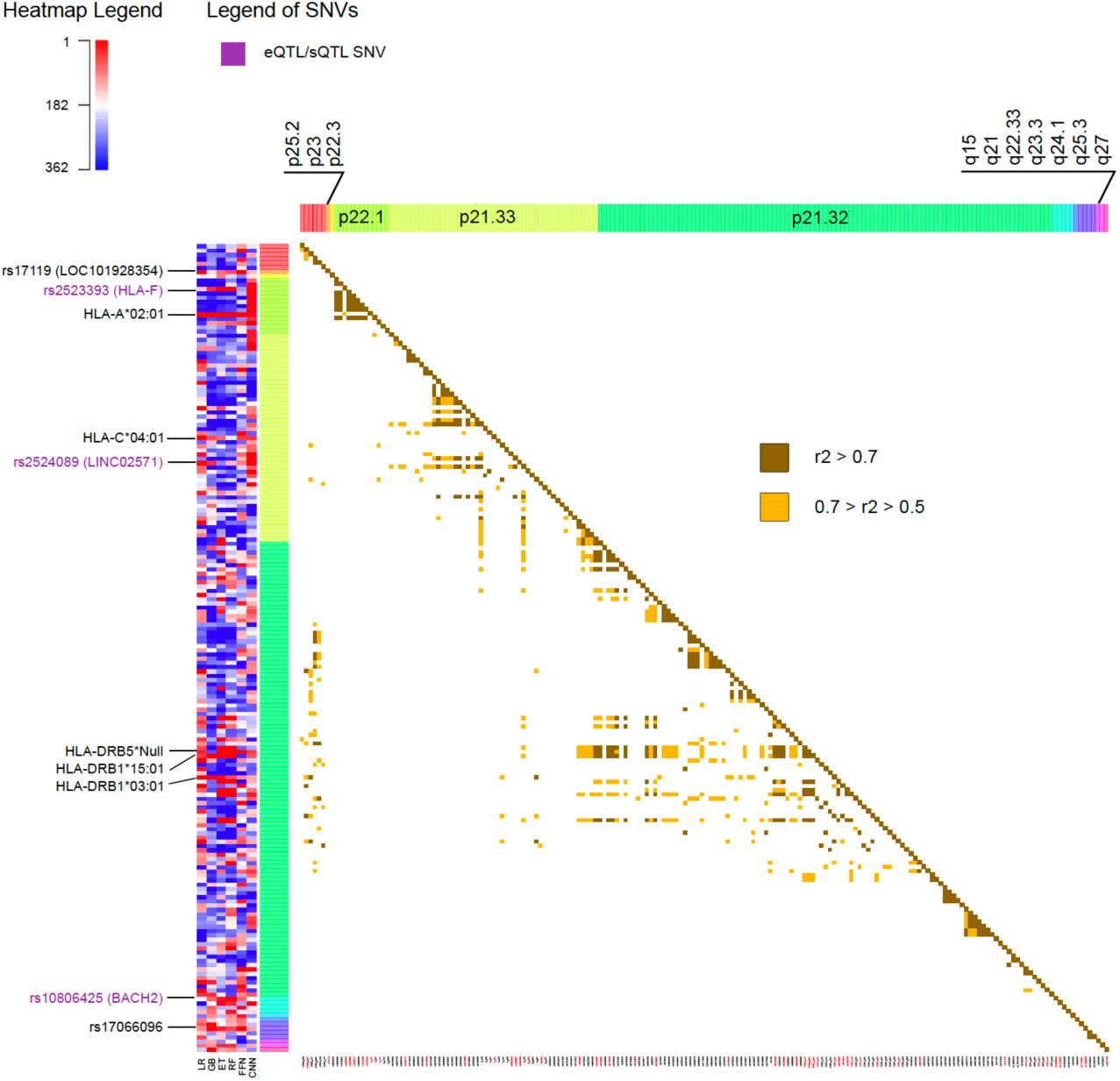
The heatmap on the left represents the ranks of all the features on chromosome 6 as assigned by each ML method, with values close to one in red indicating higher importance. The top ten best-ranked genomic variants on this chromosome are labeled with their corresponding names. Labels in purple indicate the presence of QTLs in at least one tissue in GTEx. The heatmap on the right indicates the presence and strength of LD between pairs of genomic variants.

The heatmaps in Figure 3 and Figure 4 illustrate the substantial variability of ranks assigned to genomic variants, often displaying diverse colours corresponding to ranks obtained using different ML methods. Notably, all missense variants used as features in the MS models were annotated as likely benign by AlphaMissense (Cheng et al. 2023), which aligns with the polygenic nature of MS, wherein the cumulative effect of numerous small to medium genetic effects across the genome has been described to predispose or protect against the disease (Goris et al. 2022). Alternatively, most of the SNVs were predicted to have an effect in expression (eQTL) or splicing (sQTL) as annotated using GTEx and highlighted with purple colour in the labels of the SNVs in Figure 3 and Figure 4.

Three prioritized variants were missense SNVs, highlighted with green labels in Figure 3, rs6897932 located in the *IL7R* gene of chromosome 5, rs763361 located in the *CD226* gene of chromosome 18, and rs5771069 located in the *IL17REL* gene of chromosome 22. The *IL7R* gene encodes the interleukin-7 receptor, involved in the development and function of T cells. *CD226*, on the other hand, encodes a glycoprotein also known as *DNAX accessory molecule-1* (*DNAM-1*), which plays a role in the regulation of T cell activation and the immune response. Genomic variants in *IL7R* and *CD226* have been associated with other autoimmune diseases, such as type 1 diabetes and rheumatoid arthritis (Lee et al. 2012) (Douroudis et al. 2009) (Meyer, Parmar, and Shahrara 2022) (Tan et al. 2010). Notably, there was an absence of prioritized missense variants in chromosome 6.

The top 10 ranked genomic features in chromosome 6 were determined by summing the ranks obtained with the six ML methods and are labelled in Figure 4. There is not only one location in chromosome 6 associated with MS, instead, the top ten risk loci are widely distributed across cytobands. Among the top genomic features in chromosome 6 there were five HLA types: *HLA-A*02:01*, *HLA-C*04:01*, *HLA-DRB5*Null*, *HLA-DRB1*15:01* and *HLA-DRB1*03:01*. Additionally, the SNV rs2523393, located in *HLA-F*, was identified as an eQTL and sQTL for this gene. Furthermore, the SNV rs2524089, an intron variant in *LINC02571*, was recognized as an eQTL and sQTL for the genes *HLA-B*, *HLA-C*, and *HLA-E*. In this regard, the prevalence of HLA gene annotations among the top genomic features on chromosome 6 highlights their significance in the context of MS.

The top 10 best-ranked genomic features across all chromosomes are listed in Table 5. When considering all chromosomes, the highest-ranked genomic variant was *HLA-A*02:01* on chromosome 6. In the UKB cohort, *HLA-A*02:01* was more present in controls compared to individuals with MS, with a Fisher test *p-value* of 2.43E-19. This observation aligns with its reported protective effect against MS in the literature (Brynedal et al. 2007) (Bergamaschi et al. 2010). *HLA-A*02:01* was also recurrently selected across all folds and methods with the RFE and RFECV techniques in the previous section, emphasizing the relevance of this variant for predicting MS outcomes in the UKB cohort. The *HLA-A* gene belongs to the major histocompatibility complex (MHC) class I. It is worth noting that the most significant genetic factor associated with MS, as reported in the literature, is *HLA-DRB1*15:01* (Menegatti et al. 2021), which is a predisposing HLA variant belonging to the MHC class II. In the UKB cohort, *HLA-DRB1*15:01* exhibited the most significant differences in allele frequency between individuals with MS and controls, with a Fisher test *p-value* of 2.77E-101, being more prevalent in MS than in controls, consistent with its predisposing role. However, when considering rankings across all chromosomes, this variant was ranked 23rd.

**Table 5.**
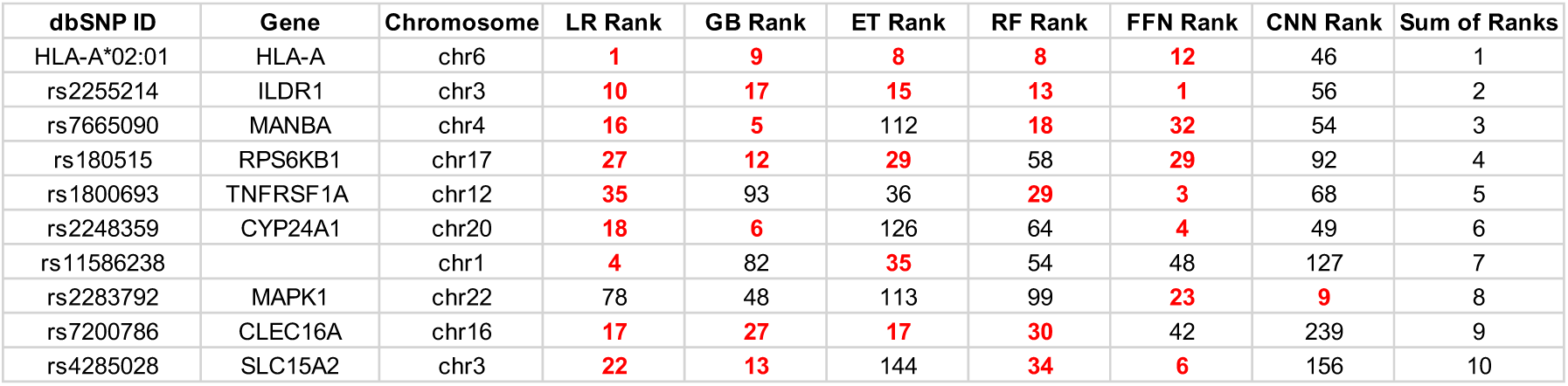
lists the top ten best-ranked genomic features across all methods with the corresponding ranks assigned by each ML method. The values of the prioritized ranks are highlighted in red.

| dbSNP ID | Gene | Chromosome | LR Rank | GB Rank | ET Rank | RF Rank | FFN Rank | CNN Rank | Sum of Ranks |
| --- | --- | --- | --- | --- | --- | --- | --- | --- | --- |
| HLA-A*02:01 | HLA-A | chr6 | 1 | 9 | 8 | 8 | 12 | 46 | 1 |
| rs2255214 | ILDR1 | chr3 | 10 | 17 | 15 | 13 | 1 | 56 | 2 |
| rs7665090 | MANBA | chr4 | 16 | 5 | 112 | 18 | 32 | 54 | 3 |
| rs180515 | RPS6KB1 | chr17 | 27 | 12 | 29 | 58 | 29 | 92 | 4 |
| rs1800693 | TNFRSF1A | chr12 | 35 | 93 | 36 | 29 | 3 | 68 | 5 |
| rs2248359 | CYP24A1 | chr20 | 18 | 6 | 126 | 64 | 4 | 49 | 6 |
| rs11586238 |  | chr1 | 4 | 82 | 35 | 54 | 48 | 127 | 7 |
| rs2283792 | MAPK1 | chr22 | 78 | 48 | 113 | 99 | 23 | 9 | 8 |
| rs7200786 | CLEC16A | chr16 | 17 | 27 | 17 | 30 | 42 | 239 | 9 |
| rs4285028 | SLC15A2 | chr3 | 22 | 13 | 144 | 34 | 6 | 156 | 10 |

The SNVs rs7665090 and rs2248359 listed in Table 5 are located downstream and upstream of the genes *MANBA* and *CYP24A1,* respectively. Specifically, rs7665090 serves as both an sQTL and eQTL for *MANBA*, which is an exoglycosidase found in the lysosome and is present in immune system pathways (González-Jiménez et al. 2022). Alternativelly, rs2248359 functions as an eQTL for *CYP24A1*, which encodes a protein involved in the catabolism of the active form of vitamin D (Law et al. 2021).

The SNV rs180515 is situated in the 3’ UTR of *RPS6KB1* gene, and is also annotated as both an eQTL and sQTL for this gene. *RPS6KB1* is actively involved in immune response pathways, particularly in the IL-4 signalling pathway, which has been associated with the progression of MS in several studies (Gadani et al. 2012), (K. Wang et al. 2018), (Kallaur et al. 2013), (Vogelaar et al. 2018).

The variants rs1800693, rs2283792, and rs7200786, are located within the intronic regions of the genes TNFRSF1A, MAPK1, and CLEC16A, respectively. The SNV rs1800693 functions as both an eQTL and sQTL for TNFRSF1A. This gene encodes a member of the TNF receptor superfamily of proteins and is known to play a role in regulating the immune system and the initiation of inflammatory reactions (Ward-Kavanagh et al. 2016), (Javor et al. 2018), (Ottoboni et al. 2013). The SNV rs2283792 serves as an eQTL for MAPK1, which is linked to MS due to its involvement in the MAPK pathways (ten Bosch et al. 2021). The SNV rs7200786 is both an eQTL and sQTL for CLEC16A, which is a direct regulator of the MHC class II pathway in antigen-presenting cells (Van Luijn et al. 2015).

Overall, most of the highest-prioritized variants were identified as eQTL or sQTL located in non-coding regions within or near genes associated with the immune response and MS. Other SNVs were prioritized by the models but were not annotated as missense variants, eQTLs, or sQTLs affecting relevant genes. This could be partially attributed to the presence of LD, which results in highly correlated genotypes for variants located close together. While this correlation among features confers similar predictive power in the models, it does not necessarily imply that each individual variant is relevant to MS.

## DISCUSSION

In this work we investigated different aspects concerning the application of ML methods for predicting complex diseases based on genomic features. It is important to acknowledge that assessing genomic predisposition to complex diseases, which do not adhere to classic Mendelian inheritance patterns, present challenges. In this regard, some ML methods have the ability to identify complex relationships in the data that the traditional statistical methods may overlook.

The performance of ML models was evaluated and compared across folds, methods, and diseases. Lower variability across folds is often desirable in ML analyses, as it suggests more stable and reliable performance. Notably, DL methods exhibited the highest variability across folds. This could be attributed to the relatively modest sample size employed in this study, posing challenges for generalization, especially when leveraging the deeper connections inherent in DL models. Related to this, several studies have suggested that less sophisticated ML methods tend to outperform DL methods when dealing with small sample sizes (Ng et al. 2020) (Dong et al. 2022). Supporting this notion is the fact that LR exhibited stable performance across folds and diseases, and was consistently positioned among the top-performing methods. LR is known for its relatively simple algorithm and ease of implementation (Goodfellow, Bengio, and Courville 2016). Nevertheless, additive regression models such as LR, by default, are designed to detect main effects, preventing them from capturing interactions between input variables. In this respect, this limitation did not seem to negatively impact the results of LR in the current work.

Another relevant aspect of using ML methods is to investigate how they compare to PRS, which is the most widely used tool in population genomics to quantify an individual’s genetic predisposition to a disease based on multiple genomic variants. In this work, we concluded that the performance of PRS was comparable to ML methods. Consequently, the question arises of whether to select one method over the other.

ML offers several advantages over PRS. For example, PRS are limited to capturing only linear relations with the disease, as its core algorithm is a linear additive model. In contrast, ML methods, with the exception of LR, can capture complex interactions and nonlinearities. This ability could be especially valuable for detecting synergisms between genomic variants that may go unnoticed with PRS. Additionally, in the case of ML, the interpretability of the model is more flexible compared to PRS. This is because PRS provide a risk score based on a set of genomic variants with their associated weights obtained from GWAS summary statistics, and these weights remain unmodified. However, ML have the capacity to learn from the data and refine the weights assigned to genomic variants during training, following different architectures depending on the method used. Also, ML methods can produce a variety of outputs including the predicted class for each individual and its associated probability. As probabilities are easy to interpret and range from 0 to 1, the classification of the same individual can be directly compared across methods and experiments. In contrast, PRS themselves are not classifiers, and their output is a single numerical score for each individual, representing the cumulative genetic risk for a specific trait or disease. However, this score alone cannot be compared across different PRS studies, and the risk of individuals developing the disease is typically interpreted using quantiles, with the highest PRS values indicating higher genetic predisposition to the disease (Collister, Liu, and Clifton 2022).

Conversely, PRS offers several advantages over ML. One of them is that there is no need to access large datasets with individualized genomic data to build the models. This is because PRS require GWAS summary statistics instead of individualized genomic data, and summary statistics are typically anonymized to protect the privacy and confidentiality of the study participants, facilitating their public availability (Tanjo et al. 2020). Additionally, in PRS there are no limits to the number of genetic variants to include in the models, and some of the problems associated with the dimensionality of the data in ML are resolved in PRS. For example, the computation times of PRS will generally be reasonable regardless of the number of genomic variants used for the calculations due to its relatively simple core algorithm. Finally, PRS fell within the average performance when compared across methods and diseases. In contrast, the performance of ML methods was more variable, especially across tree-based ML and DL methods. It is worth noting that, apart from the methods used in this work, there is an extensive list of other ML methods that could be employed as classifiers, but it was not feasible to test all of them in this work. The uncertainty regarding the best ML method to use to solve a particular problem creates the necessity of testing several methods in the same study, adding complexity to the process of analysing the data. In this regard, PRS represents a safe choice, without the requirement of testing many different methods.

In this study we employed two recursive feature elimination tools, RFE and RFECV, with the objective of testing whether there exists a subset of predictors that could enhance the model’s performance. After applying RFE and RFECV, the number of features in the models decreased to varying extents, along with a reduction in the number of correlated features. Interestingly, there were no significant changes in the performance of the models after feature selection. The absence of performance improvement, despite the reduction in correlated features, suggests that the LD present among genomic variants did not have a major impact on the performance of the original models. The robustness towards LD was particularly unexpected for LR, which may be negatively affected by multicollinearity (Vatcheva et al. 2016).

In addition, the models developed for AD predominantly relied on a single SNV, namely rs429358, for making predictions. After the application of RFECV techniques, some models exclusively depended on this genomic variant for classification, with no significant impact on performance. Located on chromosome 19 in the *APOE* gene, the allele (C) of rs429358 is one of the most extensively reported factors associated with AD risk and dementia, exhibiting an additive risk pattern (Huang et al. 2017). In this regard, the recurrent selection of rs429358 across methods and folds, which also has substantial supporting evidence of an association with AD risk in the literature, over other variants in strong LD such as rs4420638 (*r^2^*=0.708) and rs769449 (*r^2^*=0.743) with less disease evidence, underscores the capability of RFE and RFECV to discern genomic variants with the most significant links to the disease, despite the presence of feature correlations.

The strong association between rs429358 (C) in *APOE* and the disease might overshadow the contributions of other weaker genetic risk factors in the AD models. Related to this, when trying to account for additional genomic variants conferring small risk effects to AD, it is a common practice to exclude the *APOE* region from GWAS and PRS calculations, treating the *APOE* locus as an independent factor or covariate (Bellenguez et al. 2022) (Schwartzentruber et al. 2021) (Ware et al. 2020). The same approach could be applied to ML. To do so, ensuring enough representation of the three genetic isoforms of ApoE would be desirable.

In contrast, for MS, there was a recurrent selection of the HLA variant *HLA-A*02:01* across all folds and methods with RFE and RFECV. However, none of the models entirely depended on this variant for predictions, as was the case with rs429358 in AD, supporting the presence of polygenicity in MS. Consequently, we explored the ranking of genomic features assigned by different ML models for MS. The diversity in the ranking of features across methods and folds highlighted the complexity of polygenic diseases. Additionally, the presence of LD likely contributed to the instability of the importance assigned to the genomic predictors (Toloşi and Lengauer 2011). In this context, extracting general rules, such as a unique prioritization of features that applies to all methods, becomes challenging.

Generally, the top genomic variants ranked by the ML models were located in non-coding regions near genes involved in the immune response or associated with MS. In addition, most of these variants were annotated in GTEx as eQTLs or sQTLs to these genes in at least one tissue. There was an enrichment of HLA gene annotations among the prioritized genomic variants in chromosome 6. The HLA variant *HLA-DRB1*15:01,* belonging to MHC class II genes, is the strongest genetic determinant of MS as defined in the literature (Menegatti et al. 2021) (Caillier et al. 2008) (Briggs and Sept 2021) (Misicka et al. 2022). However, in our work, the most consistently prioritized genomic variant in MS was *HLA-A*02:01*, which belongs to the MHC class I. Notably, there is evidence of the independent association of *HLA-A*02:01* and *HLA-DRB1*15:01* with MS (Sawcer et al. 2011) (Brynedal et al. 2007) (Bergamaschi et al. 2010), with the former considered protective and the latter predisposing to the disease, aligning with the results of our work.

In the case of MS, the well-known region strongly associated with the disease in the vicinity of *HLA-DRB1*15:01* on chromosome 6 did not seem to overshadow the relevance of other variants located on different chromosomes. Consistent with this observation is the fact that, when exploring the genomic variants with the top 10 sum of ranks, only *HLA-A*02:01* belonged to chromosome 6, and it was not in strong LD with *HLA-DRB1*15:01*. Genomic variants near genes such as *MAPK1* in chromosome 22, *CYP24A1* in chromosome 20, *RPS6KB1* in chromosome 17, *CLEC16A* in chromosome 16, *TNFRSF1A* in chromosome 12 and *MANBA* in chromosome 4 were among the top 10 prioritized genomic variants, highlighting their distribution across different chromosomes.

It is important to note that modifying any of the parameters in this study, such as employing different ML methods, selecting different genomic features, or applying ML methods to other diseases, could potentially alter some of the conclusions drawn in the current study. Indeed, one of the weaknesses of the ML methods observed in this work is their variability when certain conditions in the analysis are changed. Despite this limitation, ML methods have proven to be powerful and efficient in various everyday applications. In this regard, with the reduction in genome sequencing costs and improvements in sequencing technologies, the volume of genomic data is expected to continue increasing in the coming years. Simultaneously, the rise in computational capacities and advancements in existing ML methods are likely to increase the robustness of these methods and foster exciting discoveries in the field of population genomics.

## METHODS

### Inclusion and exclusion criteria

UK Biobank (UKB) was the main source of genomic data for this work. Additionally, the study conducted by IMSGC (Hafler et al. 2007) and available in dbGAP under the accession ID “phs000139.v1.p1” was used as external validation dataset for MS. ADNI was used as an external validation dataset for AD. The inclusion and exclusion criteria used to select cases and controls in each dataset is described in the supplementary material. The number of individuals from UKB, IMSGC and ADNI employed in this study, across diseases and genders, after applying the selection criteria is also provided in supplementary material.

### Pre-processing of genomic data

UK Biobank Axiom Array was the source of genomic data. From all the genomic variants present in the array, only those that were reported in ClinVar (Landrum et al. 2018) with at least one level of review status or reported in DisGeNet (Piñero et al. 2017) within the curated dataset associated with the diseases under study were used as predictors in the ML models. A binary feature indicating sex was also included in the models. When an HLA gene was associated with the disease, the imputed HLA types for this gene obtained from UKB (UKB Field ID 22182) were included as predictors. The numbers of predictors used in each disease are shown in supplementary material and the list of dbSNP IDs used as predictors for each disease is provided in supplementary tables.

Genetic variants were encoded as 0, 1, 2 and 3 corresponding to missing value, the absence of the variant, the presence of the variant in one allele, and the presence of the variant in two alleles, respectively, assuming an additive model. Genomic variants with the same values in all samples (monomorphic predictors) were excluded from the analysis. PLINK (Purcell et al. 2007) was used for the quality control. SNVs with a Hardy-Weinberg equilibrium *p-value* lower than 1e-8, minor allele frequency lower than 0.05, missingness per marker higher than 0.2, and samples with missingness per individual higher than 0.2, were excluded. Additionally, PLINK was employed to compute the linkage disequilibrium (LD) statistics between genomic variants.

The pipeline for imputing missing values involved several steps, and only the genomic variants that were already present in the UK Biobank Axiom Array but had missing genotypes in some samples (less than 20% of samples after QC filters) were imputed. Haplotype phasing was performed using SHAPEIT4 (Delaneau et al. 2019), while IMPUTE5 (Rubinacci, Delaneau, and Marchini 2020) was employed for genomic imputation. The reference files for genomic imputation were obtained from the 1000 genomes phase3 (Auton et al. 2015). Imputed genotypes with less than 80% probability were considered as missing, and imputed genomic variants with a quality score lower than 0.90 were excluded from further analysis.

### Machine learning methods

Nested cross-validation (nested CV) was applied with 10 folds in the inner loop and 5 folds in the outer loop to select the optimum hyperparameter configuration and obtain an estimate of the model’s generalization performance. For the hyperparameter selection, the grid search approach was employed, and the 10 evaluation scores obtained for each hyperparameter configuration in the inner loop were used to select the optimum hyperparameter configuration. The hyperparameter configurations were ranked in decreasing order using the mean of balanced accuracy across the 10 inner folds. From the top 10 hyperparameter configurations with higher values of balanced accuracy mean, the hyperparameter configuration with the highest value of sensitivity minus the standard deviation of sensitivity across the 10 folds was selected. For each fold in the outer loop, the selected hyperparameter configuration in the inner loop was applied in the outer loop using 80% of balanced samples for training and 20% of samples for testing. The hyperparameters considered in the grid search, the architecture of FFN and CNN with the list of fixed and tuned parameters, and the final hyperparameter configurations selected for each fold, ML method and disease are provided in supplementary material. For each ML method, evaluation metrics are presented with the mean and standard deviation across the 5 final models obtained from outer loop of the nested cross-validation (5-fold CV).

Recursive feature elimination (RFE) was implemented using the sklearn.feature_selection.RFE function in Python. Different number of features were tested using the sklearn.model_selection.GridSearchCV function, with 20, 50, 100, 150, 200, and 250 for MS, and 5, 20, 50, 100 and 150 for AD. Additionally, recursive feature elimination with cross-validation (RFECV) was implemented using the sklearn.feature_selection.RFECV function. In RFE and RFECV, balanced accuracy was used as the scoring function.

### Polygenic risk score

PRSice-2 was used to calculate the PRS (Choi and O’Reilly 2019). PLINK files from UKB were used as target data. The GWAS summary statistics used as the base data were downloaded from the NHGRI-EBI GWAS Catalog (Sollis et al. 2023) on 25/05/2023 for the studies GCST005531 (Beecham et al. 2013) related to MS and GCST007511 (Kunkle et al. 2019) to AD. The sex variable and the first 10 principal components (PC) available for researchers to download from UKB (UKB field ID 22009) were added as covariates in the PRS models. PRS were calculated five times for each disease, including in the regression model the same samples used in the folds of the outer loop of the nested CV used for training the final ML models. All the genomic variants present in the target and base data passing the quality control were used for the PRS calculation, and the number of features in the PRS models after C+T is provided in supplementary tables. Relative risk (RR) and odds ratio (OR) were used to evaluate the models, with the formulas provided below:

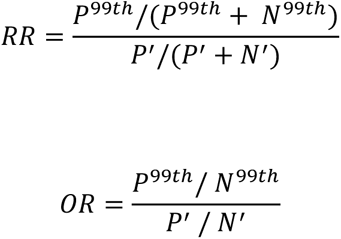

Where P^99th^ and N^99th^ represent the number of positives (individuals with the disease) and negatives (controls) present in the top 99^th^ percentile with the highest PRS, or probabilities in the case of ML methods. P^’^ and N^’^ represent the number of positives and negatives present in the samples that were not in in the top 99^th^ percentile.

### Explainability methods applied to ML models

For the tree-based ensemble ML methods such as GB, ET and RF, the importance measures assigned to the features in the classification were obtained from the feature importance metrics. In the case of LR, the coefficients of the features in the decision function were used to retrieve the importance of the features as assigned by the models. In DL methods, specifically FFN and CNN, importance metrics were derived using layer integrated gradients (LIG) (Sundararajan, Taly, and Yan 2017). To address the fact that the importance measures were obtained using different approaches, and consequently, had a different range of values, the predictors were ranked from the highest importance to the lowest importance using consecutive increasing ordinal numbers for each method and fold. The prioritization of genomic features was performed in the fold with the highest balanced accuracy for each ML method. The top 10% of the best-ranked features obtained with each ML method were selected as the prioritized genomic variants, indicating a stronger association with the disease.

## DATA ACCESS

The data used in this study were obtained from UK Biobank, dbGaP, and ADNI repositories under controlled access.

## ACKNOWLEDGMENTS

This research has been conducted using the UK Biobank Resource under Application Number 79450. We acknowledge the work from (Hafler et al. 2007) and the International Multiple Sclerosis Genetics Consortium (IMSGC) consortium for providing us with access to the dbGaP dataset with the accession number phs000139.v1.p1. Data collection and sharing for this project was funded by the Alzheimer’s Disease Neuroimaging Initiative (ADNI) (National Institutes of Health Grant U01 AG024904) and DOD ADNI (Department of Defense award number W81XWH-12-2-0012). ADNI is funded by the National Institute on Aging, the National Institute of Biomedical Imaging and Bioengineering, and through generous contributions from the following: AbbVie, Alzheimer’s Association; Alzheimer’s Drug Discovery Foundation; Araclon Biotech; BioClinica, Inc.; Biogen; Bristol-Myers Squibb Company; CereSpir, Inc.; Cogstate; Eisai Inc.; Elan Pharmaceuticals, Inc.; Eli Lilly and Company; EuroImmun; F. Hoffmann-La Roche Ltd and its affiliated company Genentech, Inc.; Fujirebio; GE Healthcare; IXICO Ltd.; Janssen Alzheimer Immunotherapy Research & Development, LLC.; Johnson & Johnson Pharmaceutical Research & Development LLC.; Lumosity; Lundbeck; Merck & Co., Inc.; Meso Scale Diagnostics, LLC.; NeuroRx Research; Neurotrack Technologies; Novartis Pharmaceuticals Corporation; Pfizer Inc.; Piramal Imaging; Servier; Takeda Pharmaceutical Company; and Transition Therapeutics. The Canadian Institutes of Health Research is providing funds to support ADNI clinical sites in Canada. Private sector contributions are facilitated by the Foundation for the National Institutes of Health (www.fnih.org). The grantee organization is the Northern California Institute for Research and Education, and the study is coordinated by the Alzheimer’s Therapeutic Research Institute at the University of Southern California. ADNI data are disseminated by the Laboratory for Neuro Imaging at the University of Southern California.

