## Supplementary material for "Machine learning methods applied to classify complex diseases using genomic data"

### Index

#### Supplementary Figures

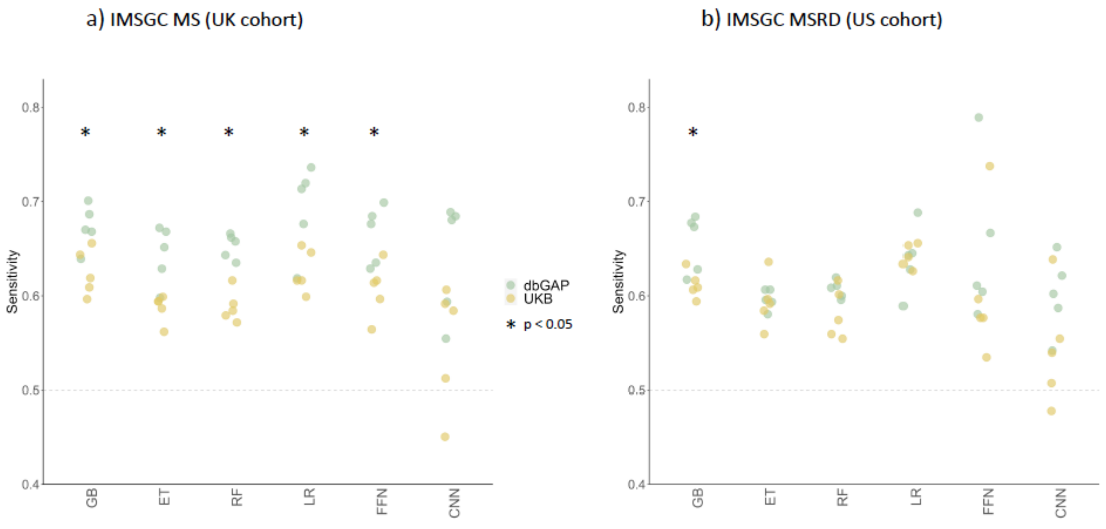

*Figure 1 shows the sensitivity values of the five folds in the outer loop resulting from training the models with the UKB cohort, and testing the models in the dbGAP (in green) and UKB (in yellow) cohorts. In (a), the results are shown for the IMSGC MS cohort, and in (b), for the IMSGC MSRD cohort. The significance of the differences between IMSGC and UKB cohorts was assessed using a Wilcoxon rank-sum test.*

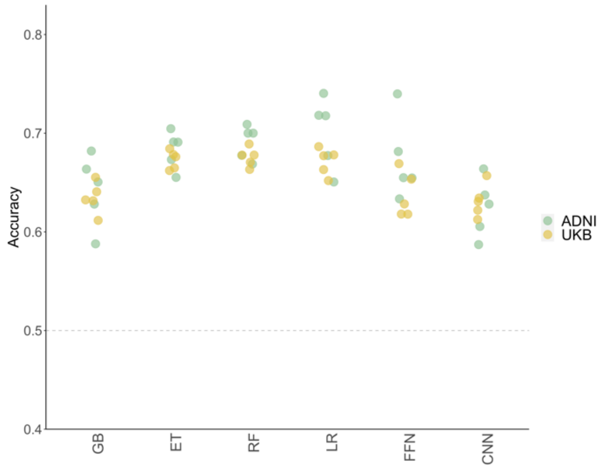

*Figure 2 shows the values of balanced accuracy in the five folds of the outer loop, resulting from training the models with the UKB cohort, and testing the models in the ADNI (in green) and UKB (in yellow) cohorts.*

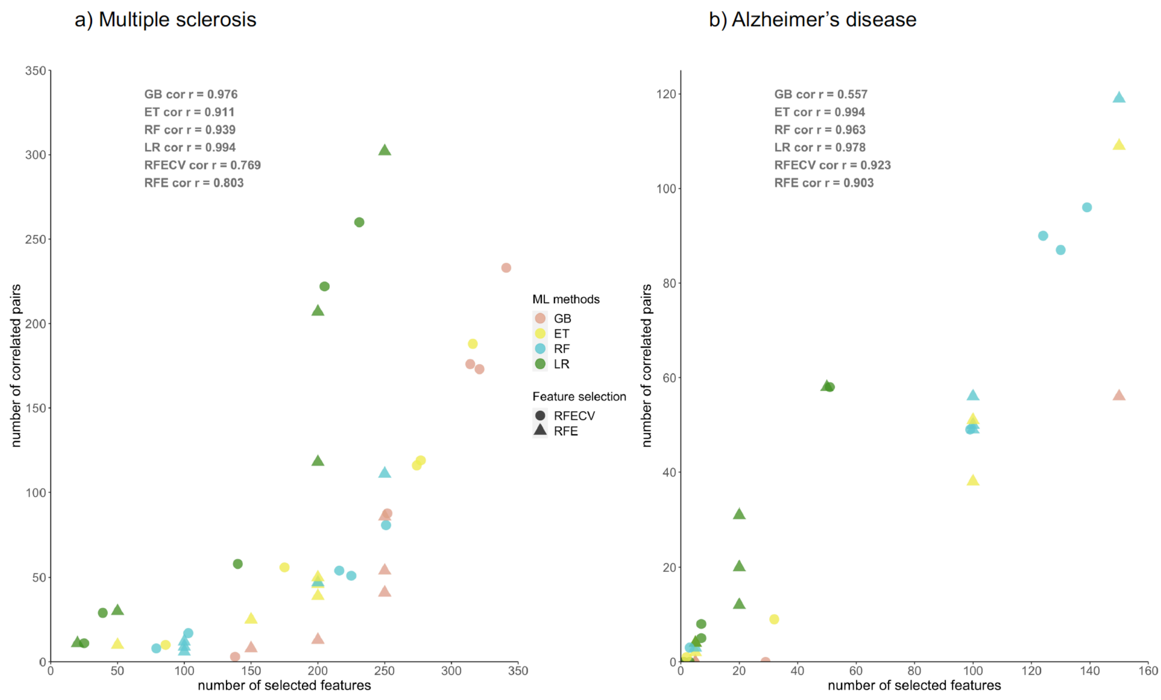

*Figure 3 displays the relationship between the number of selected features and the number of correlated pairs of features after the application of feature selection tools. The plots are divided into MS and AD represented in (a) and (b) respectively. The correlation coefficients were calculated using Spearman correlation.*

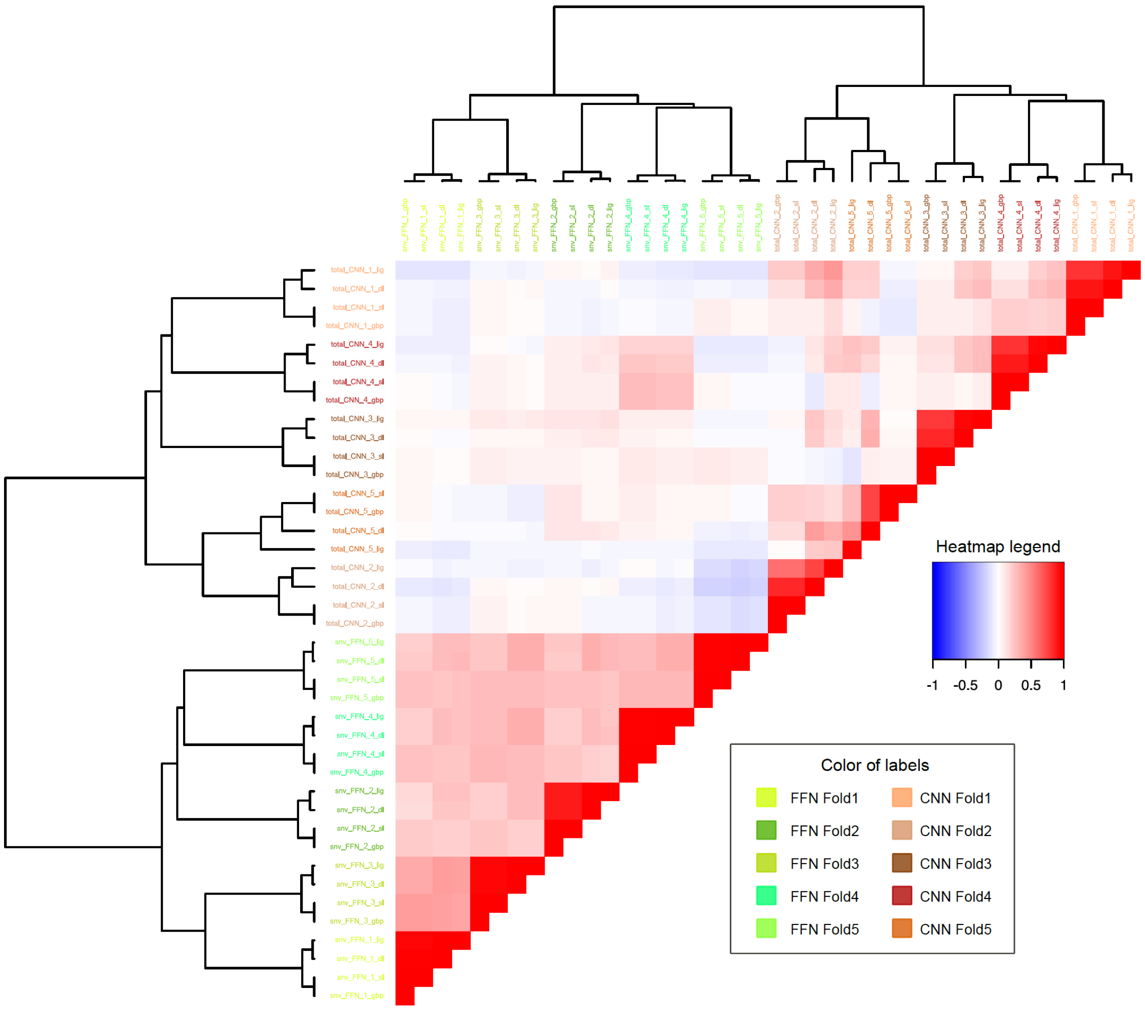

*Figure 4 shows the pairwise Pearson correlation coefficients of the ranking of features obtained with layer integrated gradients (LIG), layer deeplift (DE), saliency maps (SM) and guided backpropagation (GBP) applied to CNN depicted with warm colors in labels, and FFN depicted with green colors in labels. The clustering in the dendrogram is made with Euclidean distances and ward-D2.*

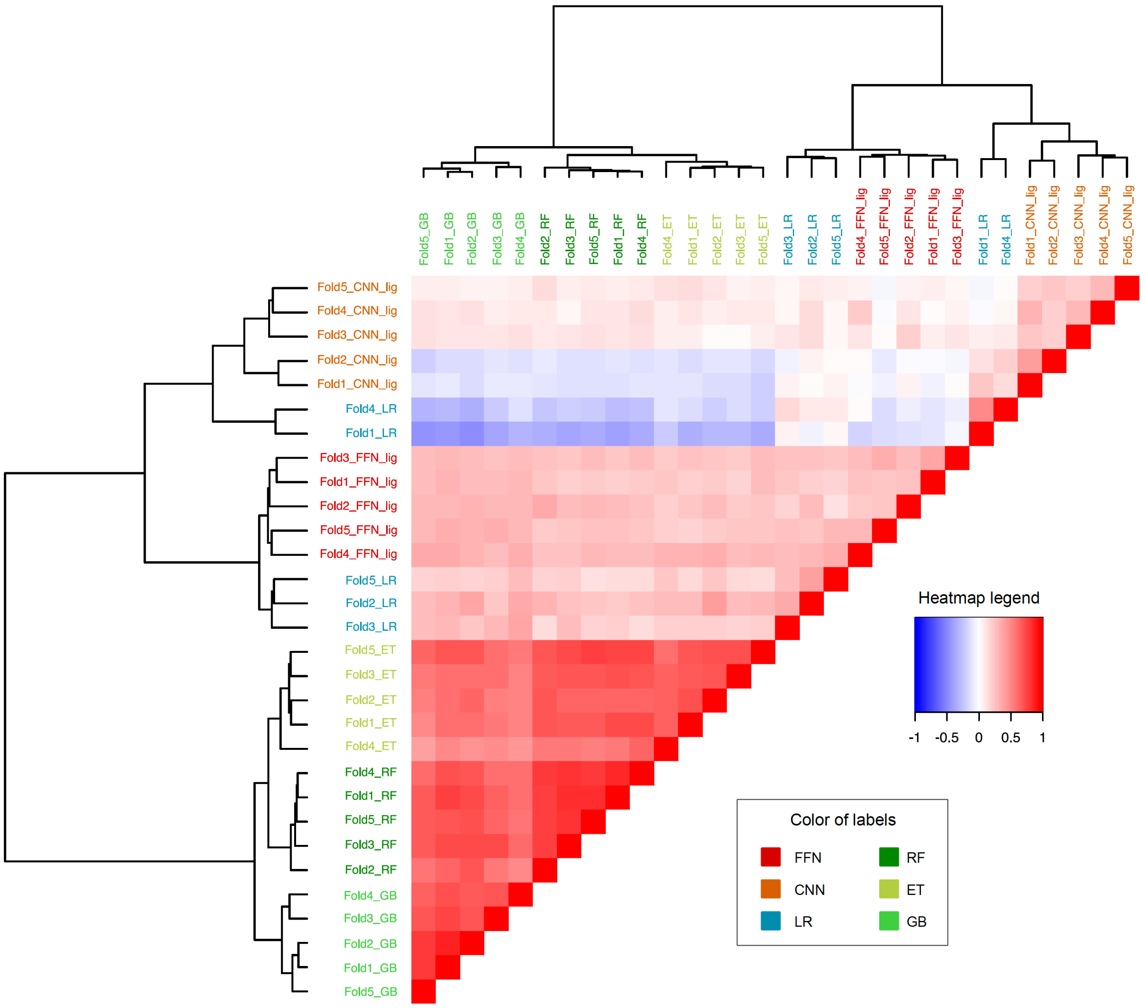

*Figure 5 shows the pairwise Pearson correlation coefficients of the ranking of features obtained with different ML methods. LIG is used for DL methods. The clustering in the dendrogram is made with Euclidean distances and ward-D2.*

#### Extended Methods

##### Inclusion and exclusion criteria

For this work, we used information from individuals in the UK Biobank (UKB). The UK Biobank Axiom Array[1], a custom-designed array manufactured by Affymetrix, was the source of genomic data. This array contains nearly 820,000 genetic markers, including SNVs, and small insertion-deletion polymorphisms (indels).

The inclusion criteria used to select cases and controls in UKB were as follows:

- MS: Subjects with the ICD-10 code G35 in primary care data, hospital inpatient data, or mortality data.
- AD: Subjects with the ICD-10 code G30.9 in primary care data, hospital inpatient data, or mortality data. 78 subjects (3% of the total AD) had less than 65 years old and therefore, were probable EOAD. We decided to include the probable EOAD in the analysis under the premise that the models may be able to correctly classify them as AD, as both EOAD and LOAD share some genetic determinants. In any case, the age at first report was explored to evaluate potential biases in age among the true positives and false negatives as classified by the models (Figure 6).
- Controls: Subjects who are more than 75 years old and do not have any disease of the nervous system or mental, behavioral, or neurodevelopmental disorders (ICD-10 categories G00-G99 and F01-F99).

The exclusion criteria applied to subjects in UKB were as follows:

- Subjects without any clinical information.
- Subjects with more than one of the studied diseases.
- Genetic ethnic grouping not Caucasian (UKB Field ID 22006).
- Recommended genomic analysis exclusions due to poor heterozygosity/missingness (UKB Field ID 22010).

- Individuals with high heterozygosity rate (after correcting for ancestry) or high missing rates (UKB field ID 22018).
- Individuals with sex chromosome aneuploidy (UKB field ID 22019).
- Outliers for heterozygosity or missing rate (UKB field ID 22027).
- From the genetically related individuals, only one subject (preferentially with the disease) was included in the analysis (UKB field ID 22011).

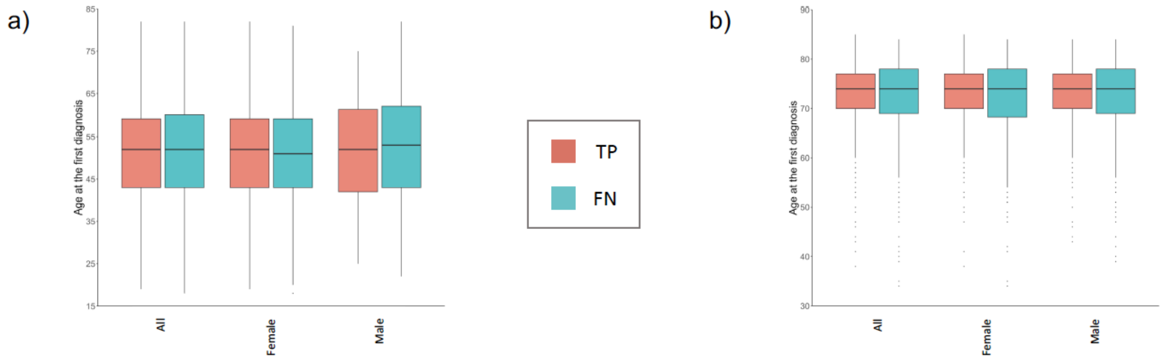

*Figure 6 illustrates the differences in the age of individuals at the first report of the disease in the UKB clinical records. TP corresponds to individuals with the disease who were correctly classified as having the disease by all ML methods, while FN corresponds to individuals with the disease who were incorrectly classified by at least one method. The significance of the differences between TP and FN was assessed using a t-test. The plots in (a) and (b) represent the results for MS and AD, respectively.*

| | **Cases** | | **Controls** | |
|---|---|---|---|---|
| | **Female** | **Male** | **Female** | **Male** |
| **MS** | 1443 | 577 | 42154 | 37695 |
| **AD** | 1357 | 1133 | 42154 | 37691 |

*Table 1 shows the distribution of individuals from UKB employed in this study, across diseases and genders, after applying the selection criteria.*

The ADNI dataset (adni.loni.usc.edu)[2] was used as an external validation dataset for AD. In this study the data coming from whole-genome sequence (WGS) at high coverage was used. The inclusion criteria used to select cases and controls in ADNI were as follows:

- AD: Individuals with probable or possible diagnosis of AD (field name: DXAPP) and dementia due to AD (field name: DXDDUE) or mild cognitive impairement (MCI) due to AD (field name: DXMDUE).
- Controls: Individuals without MCI or dementia (field name: DIAGNOSIS), without probable or possible diagnosis of AD (field name: DXAPP), without dementia due to AD (field name: DXDDUE) and without MCI due to AD (field name: DXMDUE).

The exclusion criteria applied to subjects in ADNI were as follows:

- Individuals without WGS data available.
- Individuals with missing data in the demographic variables gender and year of birth (field names: PTGENDER and PTDOBYY).
- Individuals with missing values in the examination date (field name: EXAMDATE).
- Individuals with less than 75 years old.

| | Cases | | Controls | |
|---|---|---|---|---|
| | Female | Male | Female | Male |
| ADNI | 17 | 39 | 17 | 20 |

*Table 2 shows the distribution of individuals in the ADNI dataset after applying the selection criteria.*

Data from Affymetrix GeneChip® Human Mapping 500K arrays generated by the International Multiple Sclerosis Genetics Consortium (IMSGC) and available in dbGAP under the accession ID "phs000139.v1.p1" was used as external validation dataset for MS. The dataset consisted in two cohorts: named as "multiple sclerosis" (IMSGC MS), which included trio families recruited from across the UK, and "multiple sclerosis and related disorders" (IMSGC MSRD), comprising trio families recruited from across the US. Approximately 4% of the subjects in the latter cohort were diagnosed with clinically isolated syndrome (CIS) at the time of enrolment into the study. Additional information regarding the selection of participants can be found in the supplementary appendix of the original paper[3]. The inclusion criteria used to select cases in the IMSGC dataset were as follows:

- Subjects with MS (variable name: AFFECTION_STATUS).

- Only one individual per family was included.

The exclusion criteria applied to subjects in the IMSGC dataset were as follows:

- The data consisted in family trios. Consequently, controls were excluded from the analysis as they had at least one relative with MS.
- Subjects with more than 20% of missing genotypes were also excluded.

| | Cases | |
|---|---|---|
| | **Female** | **Male** |
| **IMSGC MS** | 363 | 122 |
| **IMSGC MSRD** | 357 | 108 |

*Table 3 shows the distribution of individuals in the IMSGC dataset after applying the selection criteria. In the case of IMSGC, two cohorts were available, IMSGC MS corresponding to individuals from the UK, and IMSGC MSRD corresponding to individuals from the US.*

**Pre-processing of genomic data**

ML methods were employed to classify cases and controls using a set of genomic variants as features in the models. These genomic variants, were reported in ClinVar[4] with at least one level of review status or reported in DisGeNet[5] within the curated dataset. A binary feature indicating sex was also included in the models. When an HLA gene was associated with the disease, the imputed HLA variants for this gene obtained from UKB (UKB Field ID 22182) were included as predictors. The numbers of predictors used in each disease are shown in Table 4.

Genetic variants were encoded as 0, 1, 2 and 3 corresponding to missing value, the absence of the variant, the presence of the variant in one copy, and the presence of the variant in two copies, respectively, assuming an additive model. Genomic variants with the same values in all samples (monomorphic predictors) were excluded from the analysis.

PLINK[6] was used to apply an initial quality control and to pre-process the genomic raw data. SNVs with a Hardy-Weinberg equilibrium *p-value* ("HWE" in PLINK) lower than 1e-8, minor allele frequency ("MAF" in PLINK) lower than 0.05, missingness per marker ("geno" in PLINK) higher than 0.2, and samples with missingness per individual ("mind" in PLINK) higher than 0.2, were excluded. Additionally, PLINK was employed to compute the linkage disequilibrium (LD) statistics between genomic variants.

| | MS | AD |
|---|---|---|
| **Features by type** | 309 SNVs<br>53 HLA<br>1 sex | 167 SNVs<br>2 HLA<br>1 sex |
| **Total features** | 363 | 170 |

*Table 4 indicates the number of features of each type used in the models for each disease.*

In genotyping arrays and WGS, missing values are not randomly distributed, and specific tools can be used for the imputation[7 8]. The pipeline for imputing missing values involved several steps, and only the genomic variants that were already present in the array but had missing genotypes in some samples (less than 20% of samples after QC filters) were imputed. The pre-processing of genomic files was performed using PLINK and bcftools[9], which included tasks such as strand flipping, genome build, and ID conversion. Haplotype phasing was performed using SHAPEIT4[10], while IMPUTE5[11] was employed for genomic imputation. The reference files for genomic imputation were obtained from the 1000 genomes phase3[12]. Imputed genotypes with less than 80% probability were considered as missing, and imputed genomic variants with a quality score lower than 0.90 were excluded from further analysis. In an attempt to impute the HLA genes, HIBAG[13] was applied to the dataset sourced from dbGAP phs000139.v1.p1 (GeneChip® Human Mapping 500K arrays). However, the quality of HLA imputation in this dataset did not meet the desired standards, and consequently, the imputed HLA types were not included in the analysis for the dbGAP dataset.

#### Machine learning models

Nested cross-validation (nested CV) was applied with 10 folds in the inner loop and 5 folds in the outer loop to select the optimum hyperparameter configuration and obtain an estimate of the model's generalization performance. For the hyperparameter selection, the grid search approach was employed, and the 10 evaluation scores obtained for each hyperparameter configuration in the inner loop were used to select the optimum hyperparameter configuration. The hyperparameter configurations were ranked in decreasing order using the mean of balanced accuracy across the 10 inner folds. From the top 10 hyperparameter configurations with higher values of balanced accuracy mean, the hyperparameter configuration with the highest value of sensitivity minus the standard deviation of sensitivity across the 10 folds was selected. For each fold in the outer loop, the selected hyperparameter configuration in the inner loop was applied in the outer loop using 80% of balanced samples for training and 20% of samples for testing. The strategy of nested CV used in this study is represented in Figure 7. The ML methods used, along with the corresponding hyperparameters considered in the grid search, are listed in Table 5.

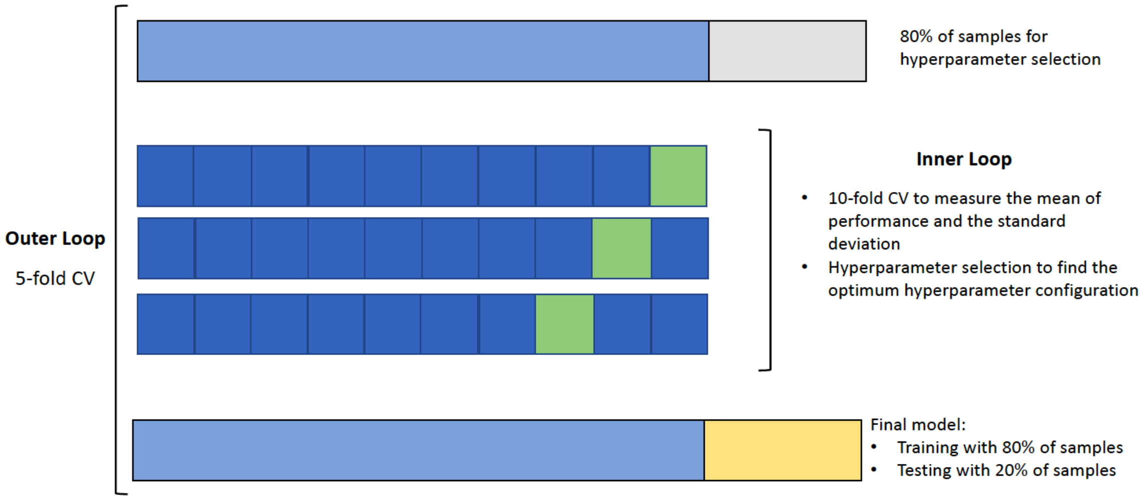

*Figure 7 represents the nested CV approach used in this work, which consists in 10-fold CV for the inner loop, and 5-fold CV for the outer loop.*

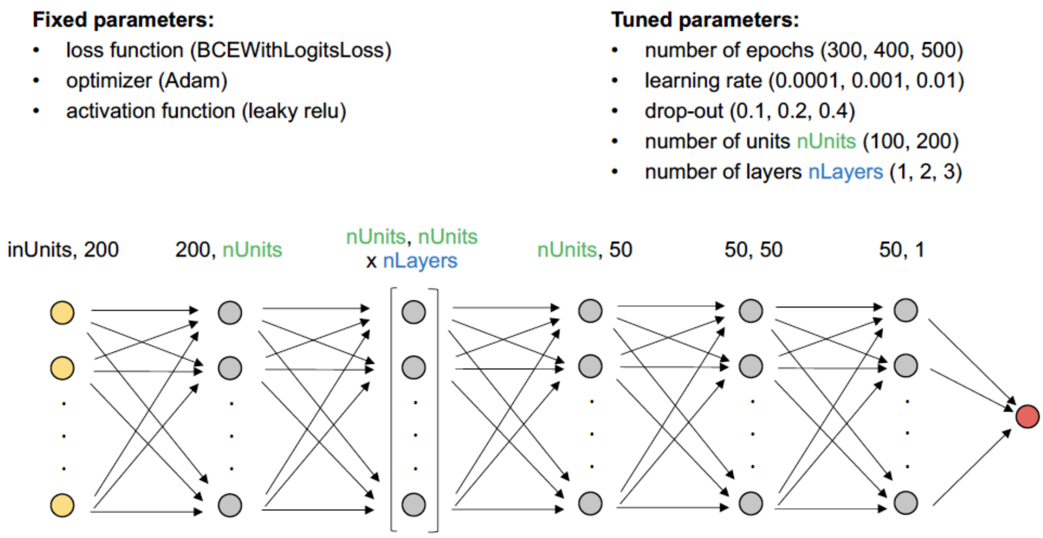

*Figure 8 represents the architecture of the FFN employed in this study.*

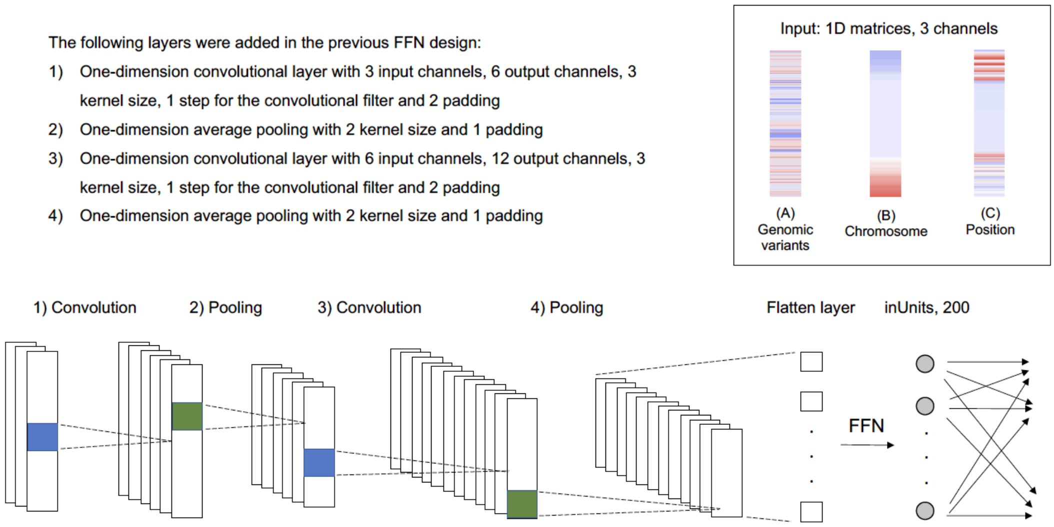

*Figure 9 illustrates the architecture of the one-dimensional CNN used in this study.*

The architecture of FFN with the list of fixed and tuned parameters used in this study is represented in Figure 8. The architecture of the CNN used in this study is represented in Figure 9. The convolutional block in the CNN employed three matrices as input channels. Matrix A represented the presence of genomic variants, like the matrices used in the other ML methods. Matrix B and matrix C represented the chromosome and position of the genomic variants, respectively. The sex variable was encoded in matrix A, with a value of 1 for females and 2 for males, while it was 0 in matrix B, and the lowest genomic position minus one in matrix C. The values in the three matrices were converted to the range of -1 to 1.

Genomic variants were ordered by chromosome and position to represent their location over the entire genome.

<table>
<tr><th></th><th>Python library</th><th>Hyperparameters</th></tr>
<tr><td>Gradient-Boosted Decision Trees (GB)</td><td>scikit-learn, GradientBoostingClassifier</td><td>• n_estimators (70, 80, 90, 100)<br>• learning_rate (0.0001, 0.001, 0.01, 0.1, 1.0)<br>• subsample (0.5, 0.7, 1.0)<br>• max_depth (7, 9, 10, 12, 14)<br>• loss ('log_loss', 'exponential')<br>• balance (50, 60, 70)<br>• sampling ('ENN', 'random', 'SMOTE_ENN', 'SMOTE_random')</td></tr>
<tr><td>Extremely Randomized Trees (ET)</td><td>scikit-learn, ExtraTreesClassifier</td><td rowspan="2">• n_estimators (50, 60, 70, 80, 100)<br>• min_samples_split (2, 5, 8)<br>• min_samples_leaf (1, 2, 5)<br>• max_depth (None)<br>• balance (50, 60, 70)<br>• sampling ('ENN', 'random', 'SMOTE_ENN', 'SMOTE_random')</td></tr>
<tr><td>Random Forest (RF)</td><td>scikit-learn, RandomForestClassifier</td></tr>
<tr><td>Logistic Regression (LR)</td><td>scikit-learn, LogisticRegression</td><td>• solver ('newton-cg', 'liblinear', 'sag', 'saga')<br>• creg (0.00001, 0.0001, 0.001, 0.01, 1, 10, 100)<br>• balance (50, 60, 70)<br>• sampling ('ENN', 'random', 'SMOTE_ENN', 'SMOTE_random')</td></tr>
<tr><td>Feedforward networks (FFN)</td><td>PyTorch</td><td rowspan="2">• number of epochs (300, 400, 500)<br>• learning rate (0.0001, 0.001, 0.01)<br>• drop-out (0.1, 0.2, 0.4)<br>• number of units nUnits (100, 200)<br>• number of layers nLayers (1, 2, 3)<br>• balance (50, 60, 70)<br>• sampling ('ENN', 'SMOTE_ENN')</td></tr>
<tr><td>Convolutional Neural Networks (CNN)</td><td>PyTorch</td></tr>
</table>

*Table 5 showing the ML methods used in this study, along with the corresponding python libraries and functions used to build the models, as well as the tested hyperparamaters.*

In addition to the hyperparameters related to the configuration of the ML methods above listed, other parameters associated with balancing and sampling strategies were tested. Varying degrees of class imbalance were used during training, where the number of cases remained constant, while the number of controls varied according to the following proportions:

- 50% cases and 50% of controls
- 40% cases and 60% controls
- 30% cases and 70% controls

As for the sampling strategies, four different approaches were tested:

- Random undersampling (random)
- Edited nearest neighbour undersampling (ENN)
- SMOTE oversampling 20% of cases + random undersampling (SMOTE_random)
- SMOTE oversampling 20% of cases + edited nearest neighbour undersampling (SMOTE_ENN)

The final hyperparameter configurations selected for each fold, method and disease are listed in Table 6 for MS and AD. The class imbalances 40%/60% and 30%/70% did not appear to confer any advantage to the model performance, as all the final hyperparameter configurations exhibited a class imbalance of 50%/50%. As for SMOTE oversampling, it was only selected in some of the hyperparameter configurations of DL methods.

The number of samples for each disease across the testing, validation, and training sets were as follows:

- MS: Testing (404 cases, 15970 controls); Validation (162 cases, 6388 controls); Training (1616 cases).
- AD: Testing (498 cases, 15969 controls); Validation (200 cases, 6387 controls); Training (1992 cases).

The number of controls in the training sets varied and depended on the imbalance rate. The final evaluation performance for each method was obtained from the outer loop in the nested CV. This was done by calculating the mean and standard deviation across the five different folds.

<table>
<tr><th></th><th>MS</th><th>AD</th></tr>
<tr><td>GB</td><td>Fold1: 100|0.01|0.5|12|exponential|50|random<br>Fold2: 70|0.001|0.5|12|deviance|50|ENN<br>Fold3: 70|0.0001|0.5|7|deviance|50|random<br>Fold4: 100|0.1|1|7|exponential|50|random<br>Fold5: 80|0.1|0.7|10|exponential|50|random</td><td>Fold1: 90|0.01|1|12|exponential|50|ENN<br>Fold2: 100|1|0.7|14|exponential|50|random<br>Fold3: 90|1|1|7|exponential|50|random<br>Fold4: 80|1|1|12|exponential|50|ENN<br>Fold5: 100|1|1|7|exponential|50|ENN</td></tr>
<tr><td>ET</td><td>Fold1: 50|5|2|None|50|ENN<br>Fold2: 50|5|1|None|50|ENN<br>Fold3: 60|5|2|None|50|ENN<br>Fold4: 50|8|1|None|50|ENN<br>Fold5: 50|2|2|None|50|random</td><td>Fold1: 60|5|1|None|50|ENN<br>Fold2: 50|8|1|None|50|random<br>Fold3: 50|5|1|None|50|ENN<br>Fold4: 60|5|1|None|50|ENN<br>Fold5: 70|5|1|None|50|random</td></tr>
<tr><td>RF</td><td>Fold1: 50|2|2|None|50|random<br>Fold2: 50|8|2|None|50|ENN<br>Fold3: 50|5|1|None|50|random<br>Fold4: 50|2|2|None|50|ENN<br>Fold5: 50|5|2|None|50|random</td><td>Fold1: 80|5|1|None|50|ENN<br>Fold2: 50|2|2|None|50|ENN<br>Fold3: 80|5|1|None|50|ENN<br>Fold4: 60|5|2|None|50|ENN<br>Fold5: 50|8|1|None|50|ENN</td></tr>
<tr><td>LR</td><td>Fold1: liblinear|1|50|random<br>Fold2: newton-cg|0.01|50|ENN<br>Fold3: saga|0.01|50|random<br>Fold4: saga|1|50|random<br>Fold5: newton-cg|0.01|50|ENN</td><td>Fold1: newton-cg|100|50|ENN<br>Fold2: newton-cg|10|50|ENN<br>Fold3: saga|100|50|ENN<br>Fold4: sag|100|50|ENN<br>Fold5: newton-cg|1|50|ENN</td></tr>
<tr><td>FFN</td><td>Fold1: 300|0.0001|0.1|100|1|ENN|50<br>Fold2: 400|0.0001|0.4|200|1|ENN|50<br>Fold3: 300|0.0001|0.2|100|1|ENN|50<br>Fold4: 500|0.01|0.2|100|1|ENN|50<br>Fold5: 300|0.0001|0.1|100|2|ENN|50</td><td>Fold1: 500|0.001|0.4|100|3|ENN|50<br>Fold2: 500|0.01|0.1|100|3|ENN|50<br>Fold3: 500|0.01|0.1|100|2|ENN|50<br>Fold4: 500|0.01|0.1|200|1|ENN|50<br>Fold5: 400|0.001|0.4|100|1|ENN|50</td></tr>
<tr><td>CNN</td><td>Fold1: 500|0.0001|0.1|100|2|SMOTE_ENN|50<br>Fold2: 500|0.0001|0.1|200|3|SMOTE_ENN|50<br>Fold3: 400|0.0001|0.1|200|1|ENN|50<br>Fold4: 500|0.0001|0.1|100|1|ENN|50<br>Fold5: 500|0.001|0.4|200|1|SMOTE_ENN|50</td><td>Fold1: 500|0.001|0.4|100|3|ENN|50<br>Fold2: 500|0.01|0.2|200|1|SMOTE_ENN|50<br>Fold3: 500|0.001|0.4|200|1|ENN|50<br>Fold4: 300|0.01|0.2|100|3|SMOTE_ENN|50<br>Fold5: 400|0.0001|0.1|100|3|SMOTE_ENN|50</td></tr>
</table>

*Table 6 lists the hyperparameters selected for the final models in MS and AD. The five folds correspond to the outer loop of the nested CV. The parameters listed for GB, in order, are: n_estimators, learning_rate, subsample, max_depth, loss, balancing, and sampling. The parameters listed for ET and RF, in order, are: n_estimators, min_samples_split, min_samples_leaf, max_depth, balancing, and sampling. The parameters listed for LR, in order, are: solver, C, balancing, and sampling. The parameters listed for FFN and CNN, in order, are: number of epochs, learning rate, dropout probability, number of units, number of layers, sampling, and balancing.*

Feature selection methods were used to identify a subset of predictors that could potentially enhance the performance of the models. Recursive feature elimination (RFE) was implemented using the sklearn.feature_selection.RFE function in Python. Different number of features were tested using the sklearn.model_selection.GridSearchCV function, with 20, 50, 100, 150, 200, and 250 for MS, and 5, 20, 50, 100 and 150 for AD. Additionally, recursive feature elimination with cross-validation (RFECV) was implemented using the sklearn.feature_selection.RFECV function. In RFE and RFECV, balanced accuracy was used as the scoring function.

#### Polygenic risk score

PRSice-2 was used to calculate the polygenic risk score (PRS)[14] for the disease of interest. PLINK files from UKB were used as target data. The summary statistics used as the base data were downloaded from the NHGRI-EBI GWAS Catalog[15] on 25/05/2023 for the studies GCST005531[16] related to MS and GCST007511[17] to AD.

PRS was calculated using the average effect size function and considering an additive model for regression. PRS calculation was combined with *p-value* thresholding using the C+T (*clumping + thresholding*) method[20]. Following this approach, PRS was calculated several times comprising SNVs with increasing GWAS *p-value* thresholds, and the most predictive PRS was used for the final PRS calculation.

Genomic variants in the base data were filtered to exclude multi-allelic SNVs. Discrepancies caused by inverted effect alleles were resolved, and each rsID was linked to a single nucleotide change. The sex variable and the first 10 principal components (PC) available for researchers to download from UKB (UKB field ID 22009) were added as covariates in the PRS models.

PRS were calculated five times for each disease, including in the regression model the same samples used in the outer loop of the nested CV used for training the final ML models. The aim was to compare the performance of PRS with ML using the same samples for fitting and evaluation in each fold. To convert PRS into binary categories, a threshold was established to distinguish the individuals with high risk to the disease. Individuals with a PRS above the 99$^{th}$ percentile were classified as high risk (positives)[21]. Similarly, the 99$^{th}$ percentile was applied to the probabilities obtained from ML models to classify high-risk individuals and to compare the results with the PRS models. The relative risk (RR) and odds ratio (OR) were used to evaluate the models, with the formulas provided below:

$$RR = \frac{P^{99th}/(P^{99th} + N^{99th})}{P'/(P' + N')}$$

$$OR = \frac{P^{99th}/\,N^{99th}}{P'\,/\,N'}$$

Where $P^{99th}$ and $N^{99th}$ represent the number of positives (individuals with the disease) and negatives (controls) present in the top 99th percentile with the highest PRS, or probabilities in the case of ML methods. $P'$ and $N'$ represent the number of positives and negatives present in the samples that were not in in the top 99th percentile.

**Explainability methods applied to machine learning models**

The importance measures assigned to the features in the classification were obtained through various approaches depending on the ML method. For the tree-based ensemble ML methods such as GB, ET and RF, feature importance metrics were obtained from model statistics. In the case of LR, the coefficients of the features in the decision function were used. In DL methods, specifically FFN and CNN, importance metrics were derived using layer integrated gradients (LIG)[22], layer deeplift (DE)[23], saliency maps (SM)[24] and guided backpropagation (GBP)[25]. These methods provide a score for each sample and feature, and the resulting matrices share the same dimensions as the input matrices. To obtain a single importance value for each feature, the median of the absolute values of attributes was calculated for cases and controls, and both values were summed for each feature. In the case of CNN, this process was repeated for each matrix, and the resulting values from the three matrices were summed. To address the fact that the importance measures were obtained using different approaches, and consequently, had a different range of values, the predictors were ranked from the highest importance to the lowest importance using consecutive ordinal numbers for each method and fold.

The prioritization of genomic features was performed in the fold with the highest balanced accuracy for each ML method. The top 10% of the best-ranked features in each method were selected as the prioritized genomic variants indicating a stronger association with the disease. To add information on the predicted pathogenic effect of missense mutations to the protein, AlphaMissense was employed[26]. To incorporate information regarding the impact of SNVs on RNA expression and splicing, data on expression quantitative trait loci (eQTL) and splicing quantitative trait loci (sQTL) from the GTEx database were used.
